# A circuit involving the lncRNA MB3 and the driver genes MYC and OTX2 inhibits apoptosis in Group 3 Medulloblastoma by regulating the TGF-β pathway via HMGN5

**DOI:** 10.1101/2024.10.21.619459

**Authors:** Grandioso Alessia, Tollis Paolo, Pellegrini Francesca Romana, Falvo Elisabetta, Palma Alessandro, Migliaccio Francesco, Belvedere Alessandro, Rea Jessica, Tisci Giada, Carissimo Annamaria, Bozzoni Irene, Trisciuoglio Daniela, Ballarino Monica, Ceci Pierpaolo, Laneve Pietro

## Abstract

**Background:** Group 3 (G3) is one of the most common, aggressive and fatal subtypes of the paediatric cerebellar tumour Medulloblastoma (MB), primarily driven by the MYC oncogene. Targeting MYC has long been challenging and this, combined with our incomplete understanding of G3 MB molecular bases, has hindered the development of effective targeted therapies. Long noncoding RNAs (lncRNAs), with their extensive oncogenic roles, cancer-specific expression, and connection to MYC biology, offer opportunities for unravelling this complexity and providing new insights and therapeutic targets.

**Methodology:** Using genome-wide, molecular and cellular assays, we characterised the activity of the MYC-dependent, anti-apoptotic lncRNA *lncMB3* in G3 MB cells.

**Results:** Through transcriptomic and interactomic analyses, we clarified *lncMB3* function and mode-of-action. *LncMB3* controls the TGF-β pathway, critically altered in G3 medulloblastomagenesis. This regulation occurs via the direct coding-noncoding RNA interaction between *lncMB3* and the mRNA for the epigenetic factor HMGN5, with both sharing targets in the TGF-β cascade. This axis converges on apoptosis through OTX2, another G3 MB driver gene, and photoreceptor lineage genes. Synergistic effects between *lncMB3* targeting and cisplatin treatment underscores the relevance of this regulatory network *in vitro*. Finally, we propose novel ferritin-based nanocarriers as efficient delivery tools for antisense oligonucleotides targeting *lncMB3*.

**Conclusions:** *LncMB3* emerges as a central node linking MYC amplification to apoptosis inhibition through a circuit involving RNA-based mechanisms, G3 MB key drivers and underexplored factors. This integrated framework deepens our understanding of G3 MB molecular underpinnings and lay the foundation for translating lncRNA research into potential applications.

## INTRODUCTION

Medulloblastoma (MB) is the most prevalent malignant paediatric central nervous system tumour, accounting for ∼20% of all childhood brain cancers [1], [2]. Although all MBs arise in the cerebellum, share histomorphological features, and exhibit of embryonic cerebellar lineage signatures [3], they show high heterogeneity. MB is classified into 4 subtypes, each with distinct biological, clinical and therapeutical implications [4]: Wingless (WNT), Sonic Hedgehog (SHH), Group 3 (G3) and Group 4 (G4). G3 and G4, together, constitute 60% of MB cases and are the most aggressive, posing treatment challenges due to poor biochemical characterisation. Deciphering the molecular mechanisms of these subgroups is critical for advancing precision medicine. G3 MB, in particular, presents a dismal prognosis, often with metastases at diagnosis and high recurrence rates [2], primarily driven by the oncogenic transcription factor (TF) MYC, characterized by gene copy number amplification or elevated expression [5]. Despite challenges in addressing canonical MYC circuits, targeting MYC and related pathways/complexes is essential in G3 MB management [6].

The discovery of novel classes of noncoding RNAs (ncRNAs) broadens our understanding of functional genome outputs [7], offering new therapeutic avenues for cancer. Among ncRNAs, the versatile long noncoding RNAs (lncRNAs) have emerged as key regulators of cancer hallmarks [8], showing promise as biomarkers and targets due to their cell-specific expression and regulatory roles. Over years, we have explored ncRNAs in various nervous system cancer models, including MB [9]. Recently, we discovered MYC-dependent lncRNAs in G3 MB, three of them potentially acting as oncogenes [10], including *lncMB3*. Here, we functionally analyse *lncMB3*, which is overexpressed the tumour conditions and exerts an anti-apoptotic action. We show that *lncMB3* modulates the TGF-β pathway, significantly altered in G3 MB [11], through a functional interaction with the mRNA for the epigenetic factor HMGN5, whose protein level is upregulated and impacts on a subset of TGF-β pathway genes also affected by *lncMB3* targeting. This regulatory axis influences apoptosis through OTX2, another crucial G3 MB driver gene [12], and its downstream cascade. This circuit relevance is underscored by the synergistic effects observed when *lncMB3* targeting is paired with cisplatin administration in G3 MB cells. Lastly, we demonstrate that recombinant human ferritin-based nanovectors, a promising platform for drug delivery in diseased conditions [13], effectively carry antisense oligonucleotides (ASOs) targeting *lncMB3* in G3 MB cells.

## MATERIALS AND METHODS

All cell lines were obtained from ATCC and cultured as previously described [10]. Evaluation of viable cell number was assessed after cell manipulations by CytoSMART Cell Counter. Gene silencing or overexpression were realized transfecting antisense LNA GapmeRs or plasmid constructs (EPB backbone-based) by Lipofectamine 2000 as per manifacturer’s instructions. Expression of specific RNAs was performed by standard qRT-PCR of digital PCR. TruSeq Stranded mRNA Library Prep Kit was used to obtain sequencing libraries from polyA+ RNA. RNA-Seq analysis were performed by conventional R package DESeq2 (Bioconductor), Gene ontology and Gene Set Enrichment Analysis. Protein expression was assessed by conventional immunoblotting, detected by ChemiDoc XRS+ Molecular Imager and quantified through the Image Lab Software. Chemotherapeutics (cisplatin, vincristine) were administered at different concentrations according to the specific experimental plan, eventually in combination with GapmeR transfection. Apoptosis was analysed by determination of Annexin V-FITC staining through flow cytometry. Data were analysed using the Flowing software. Alternatively, cell death markers were analysed by immunoblot assay. For Crosslinked Immunoprecipitation assay, cytoplasmic extracts were incubated with AGO2 antibody or IgG and coupled to ProteinG Dynabeads resin. Immunoprecipitated proteins were collected in RIPA buffer and analysed. Immunoprecipitated RNA was treated by Proteinase K before qRT-PCR . RNA Pull Down experiments were performed as described in [14] from endogenous cell extracts using separate sets of antisense biotinylated probes binding to streptavidin Magnasphere paramagnetic beads. Pull Down-Seq analysis were performed through edgeR to quantify significant genes, using a generalised linear model. RNA– RNA interaction prediction was computed using IntaRNA 3.3.2, whereas TargetScan Human database was exploited to obtain candidates microRNAs. Ribotagging and RNA Immunoprecipitation was performed as described in [15], exploiting exogenous expression of a FLAG-tagged version of the RPL22 protein for enriching the ribosomal fraction from cell lysates. HFt-HIS-PASE was obtained as a recombinant protein considering codon optimisation for high expression levels in *E. coli* . BL21 (DE3) strain. Size-Exclusion Chromatography experiments were performed using a Superose 6 gel-filtration column. The HFt-HIS-PASE protein was incubated with Fluorescein-5-Maleimide for fluorescent labelling. Electrophoresis on agarose gel of the purified ferritin-GapmeR complexes was used to demonstrate GapmeR loading of. For nucleocytoplasmic staining of fluo-HFT targeted cells, they were fixed in paraformaldehyde, permeabilised and blocked with Triton X-100/BSA/PBS. Fixed cells were incubated with Alexa Fluor™ 555 Phalloidin and DAPI solution. For Confocal microscopy experiments samples were imaged using an Olympus iX83 FluoView1200 laser scanning confocal microscope. The Fiji “Analyse Particles” tool was employed for cell counts, whereas signal co-localisation was analysed using the “JACoP” Fiji plugin. HFt-ASOs complexes were directly administered to cells as per the specific experimental plan. Oligonucleotide, probe and GapmeR sequences are listed in Table 2. For detailed technical descriptions, see “Supplementary Materials and Methods”.

## RESULTS

### Identification of *lncMB3*-dependent transcriptome in G3 MB cells

To uncover the molecular network downstream of *lncMB3* in MB, we conducted transcriptome analysis post-knockdown (KD) in the MYC-amplified D283 Med cell line where the lncRNA is highly expressed (Fig. S1A) and was first identified [10]. GTEx data (https://gtexportal.org/home/) show minimal expression of *lncMB3* in healthy tissues (Fig. S1B). KD reduced *lncMB3* expression by ∼70% (Fig. S2A) 72 hours after multi-pulse transfections of a Locked Nucleic Acid (LNA)-based ASO [16], named GapmeR #1, targeting the lncRNA 5’ region. Transcriptome data analysis showed robust gene detection (Fig. S2B, left panel), accurate transcriptomes’ clustering (Fig. S2B, right panels) and consistency of *lncMB3* KD normalised read counts with previous qRT-PCR analyses (Dataset 1).

RNA-Seq analysis (Dataset 1) detected 14.655 transcripts expressed in at least one sample and identified 2.995 differentially expressed genes (DEGs) between *lncMB3* KD and control (SCR), with FDR < 0,05 (Fig. 1A, left panel). Among these, 1.395 were downregulated and 1.600 upregulated following *lncMB3* silencing (Fig. 1A, right panel, blue and red dots). A dozen of these genes were randomly selected with padj ranging from 0,001 to 0,05 for qRT-PCR validation, confirming RNA-Seq expression trends (Fig. S2C and Dataset 1).

**Figure 1.**
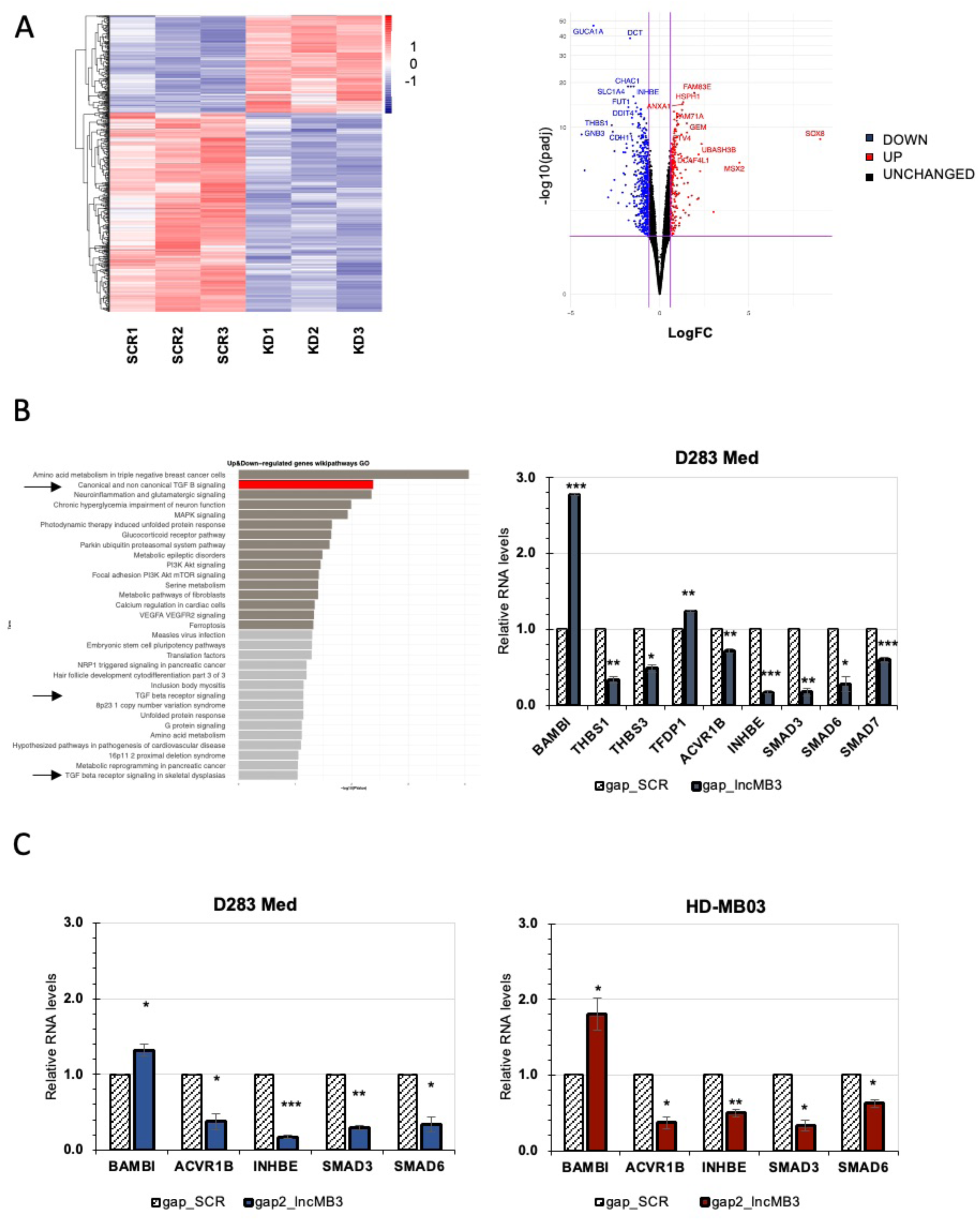
Differential gene expression analysis upon*lncMB3* KD in G3 MB cells. A. Left Panel: heatmap showing the relative levels of DEGs according to RNA-Seq analyses, along with genes hierarchical clustering (3 biological replicates for each condition). The heatmap represents only the DEGs. Right Panel: volcano plot showing DEG distribution. Genes were plotted based on statistical significance −log_10_(FDR) and differential expression log_2_(FC). Downregulated genes (logFC < -0,6) are indicated by the blue dots; upregulated genes (logFC > 0,6) by red dots. Invariant genes are indicated in black. B. GO (wikipathways) showing the distribution in clusters of the DEGs in gap_lncMB3 *vs* gap_SCR-treated D283 Med cells, according to RNA-seq data. Categories are listed considering -log_10_(pvalue) (left panel). qRT-PCR validation analysis of TGF-β pathway genes in D283 Med cells treated for 72 hours with gap_SCR or gap_*lncMB3* (GapmeR #1, right panel). Expression levels were compared to gap_SCR sample as control, set as 1. Data (means ± SEM) are expressed in arbitrary units and are relative to *GAPDH* mRNA levels. *N* = 3, * *p* ≤ 0,05, ** *p* ≤ 0,01, *** *p* ≤ 0,001 (two-tailed Student’s *t*-test). C. qRT-PCR analysis of TGF-β pathway genes in D283 Med cells treated for 72 hours with gap_SCR or gap2_*lncMB3* (GapmeR #2, left panel), or in HD-MB03 cells treated as above (GapmeR #2, right panel). Expression levels were compared to gap_SCR sample as control, set as 1. Data (means ± SEM) are expressed in arbitrary units and are relative to *GAPDH* mRNA levels. *N* = 3 to 5, * *p* ≤ 0,05, ** *p* ≤ 0,01, *** *p* ≤ 0,001 (two-tailed Student’s *t*-test).

To functionally explore the transcriptomic landscape, DEGs were filtered by quantitative criteria, such as fold change (FC) robustness, statistical significance, and expression levels (Fig. S2D). Thirteen RNAs regulated by *lncMB3* were identified (12 protein-coding and 1 lncRNA): 7 significantly downregulated and 6 upregulated (Dataset 2). qRT-PCR validated RNA-Seq data for the selected DEGs, displaying statistical significance (10 out of 13 genes) or discernible variance trends (Fig. S3A). The top gene identified was *INHBE*, a component of the TGF-β pathway [17], associated with G3 MB pathogenesis [11].

Gene Ontology (GO) analysis of DEGs highlighted the full set of functional gene categories affected by *lncMB3* depletion (Fig. 1B, left panel). Among the top ontological annotations, GO revealed pathways such as “PIK3-Akt signalling”, associated to chemosensitivity in various cancers, including MB [18], and “MAPK signalling” potentially reflecting apoptosis-induced DNA damage following *lncMB3* KD. Focus remained on the TGF-β pathway, the second most prominent gene category, deregulated in G3 compared to other MB subgroups. It appeared in both the up and downregulated gene sets (Fig. S3B), suggesting deregulation of both activatory and inhibitory elements in the same pathway. Gene Set Enrichment Analysis (GSEA) shows the significant impact of *lncMB3* KD on the TGF-β pathway in G3 MB cells (Fig. S3C).

### *LncMB3* regulates the TGF-β pathway in G3 MB

Aberrant TGF-β signalling is involved in the development of numerous diseases, including cancer [19]. In G3 MB, its dysregulation is believed to drive processes such as cell proliferation, differentiation, and survival [11].

The importance of the TGF-β cascade in our system and conditions was further highlighted by enriching the ontological category with additional DEGs involved in the pathway. We identified a set of 9 genes, including TGF-β/Activin signalling ligands (*INHBE*, [17]; *THBS1*, *THBS3*, [20]), receptors (*ACVR1B*,[21]; *BAMBI*, [22]), effectors (*SMAD3*, *SMAD6*, *SMAD7*, [21]), and downstream regulators (*TFDP1*, [23]) (Fig. S3D). Upon *lncMB3* KD, qRT-PCR showed repression of genes with activatory function in the pathway (*THBS1*, *THBS3*, *INHBE*, *ACVR1B*, *SMAD3*, *SMAD6*, *SMAD7*), with reductions ranging from 20% to 80% (Fig. 1B, right panel). Conversely, inhibitory components (*BAMBI* and *TFDP1*) were upregulated (1.5- to 3-fold, Fig. 1B, right panel), suggesting a coherent and coordinated contribution of both positive and negative effects leading to overall downregulation of the TGF-β/Activin branch of the network. To ensure specificity of targeting, we employed a second GapmeR (GapmeR #2), which recognises the 3’ end of *lncMB3* (Fig. S4A), achieving 70% transcript depletion in D283 Med cells (Fig. S4B). In HD-MB03, another G3 MB cell line where MYC and *lncMB3* are upregulated (Fig. S1A) that we used to further corroborate our findings, GapmeR #2 reduced *lncMB3* levels by 50% (Fig. S4B). Validations showed similar expression changes across all the tested condition for 5 TGF-β network genes: *BAMBI*, *ACVR1B*, *INHBE*, *SMAD3*, and *SMAD6* (Fig. 1C). Since GapmeR #1 targets the 5’ end of *lncMB3* and results in a more efficient downregulation (Fig. S4B), it was preferred for subsequent experiments. Overall, the RNA-Seq analysis clearly points to a role for *lncMB3* in the regulation of the TGF-β pathway in G3 MB cells.

### Insights into apoptosis regulation by *lncMB3*

In the developing nervous system, a relevant TGF-β cascade target is the brain-specific TF OTX2, known as a G3 MB-enriched oncogene [12], upregulated in both D283 Med and HD-MB03 cells (Fig. S1A). During retinal development, OTX2 occupies a pivotal position in a photoreceptor gene hierarchy, including basic leucine zipper TFs *NRL* and *CRX* [24], both overexpressed in G3 MB, where they promote cancer survival via the anti-apoptotic factor *BCL2L1* ([25] and Fig. S3D).

To evaluate these genes in primary MB tumours, we queried a transcriptomic dataset (GSE164677) from 59 patient-derived samples across all MB subgroups and 4 controls. *NRL*, *CRX*, and *BCL2L1* expression was significantly upregulated in G3 compared to healthy tissues or other MB subgroups. Although OTX2 upregulation in G3 MB vs. cerebellum was not statistically significant, it showed a rising trend (Fig. 2A). Figure 2B (left panel) illustrates the connections among these factors. Consistently, the photoreceptor gene program appeared in several downregulated annotations in *lncMB3* KD-dependent GSEA (Fig. 2B, middle panel).

**Figure 2.**
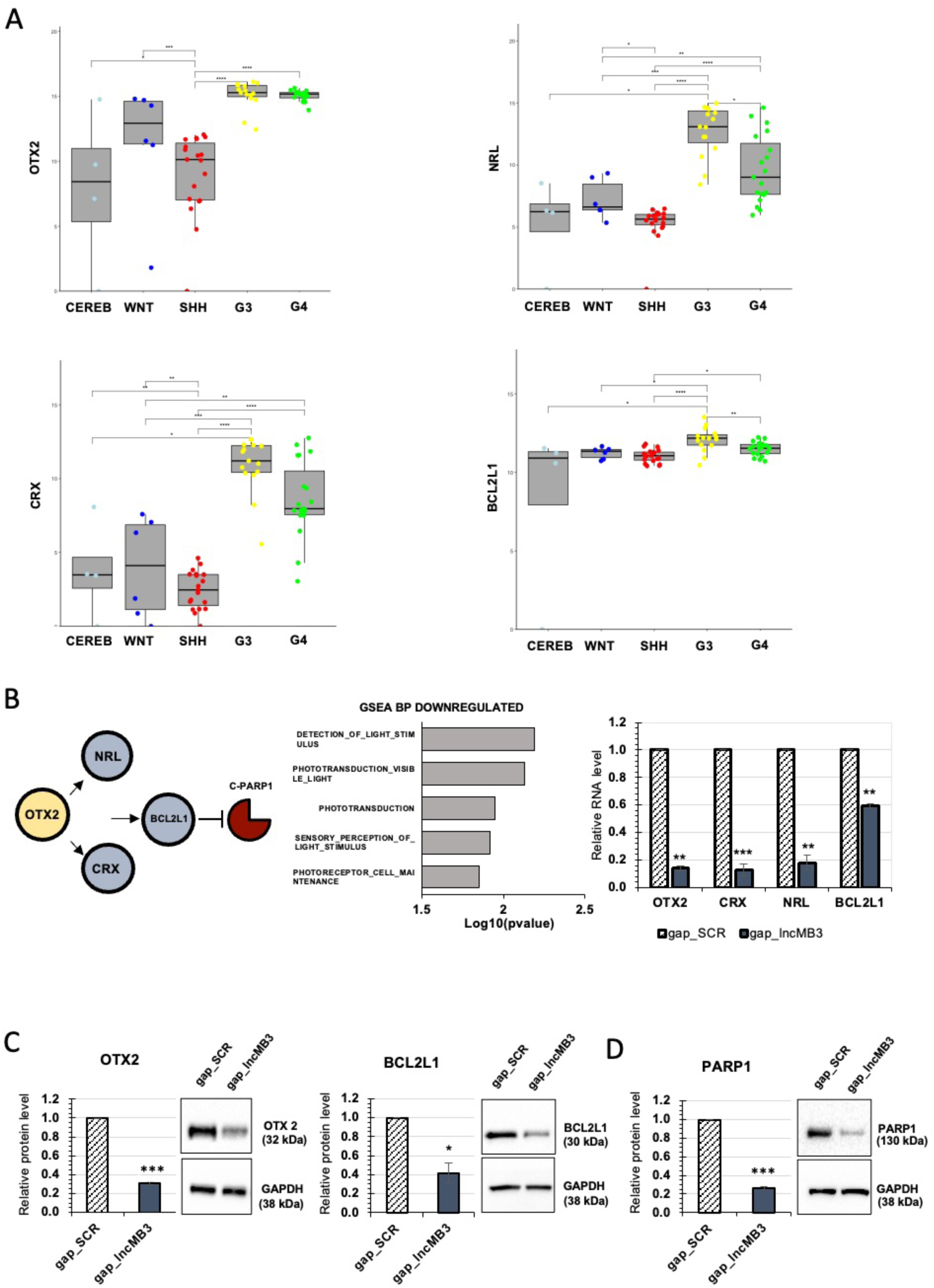
Expression of *OTX2*, *NRL*, *CRX* and *BCL2L1* in MB. A. *OTX2*, *NRL*, *CRX* and *BCL2L1* mRNA expression in 4 healthy cerebella (normal samples) and 59 primary MB samples according to the transcriptomic dataset GSE164677. Results are expressed in log_2_(FPKM). * p ≤ 0,05, ** p ≤ 0,01, *** p ≤ 0,001 **** p ≤ 0,0001 (two-tailed Student’s *t*-test). B. Schematic representation of *OTX2* network on apoptosis (left panel). GSEA biological process (BP) domains performed on deregulated genes. Categories are listed considering -log_10_(pvalue) (middle panel). qRT-PCR analysis of *OTX2*, *NRL*, *CRX* and *BCL2L1* in D283 Med cells treated for 72 hours with gap_SCR or gap_*lncMB3* (GapmeR #1, right panel). Expression levels were compared to gap_SCR sample as control, set as 1. Data (means ± SEM) are expressed in arbitrary units and are relative to *GAPDH* mRNA levels. *N* = 3, ** p ≤ 0,01, *** p ≤ 0,001 (two-tailed Student’s *t*-test). C. Western blot analysis of OTX2 and BCL2L1 protein levels in D283 Med cells treated for 72 hours with gap_SCR or gap_*lncMB3* (GapmeR #1). Normalisations were performed relative to GAPDH protein levels. N = 3, * *p* ≤ 0,05, *** *p* ≤ 0,001, (two-tailed Student’s t-test). **D.** Western blot analysis of PARP1 protein levels in D283 Med cells treated for 72 hours with gap_SCR or gap_*lncMB3* (GapmeR #1). Details as in panel C.

*OTX2*, *NRL*, *CRX*, and *BCL2L1* RNAs were downregulated in D283 Med cells after *lncMB3* depletion (Fig. 2B, right panel). Protein downregulation was also assessed for OTX2 and BCL2L1 (Fig. 2C), as well as for PARP-1 (Fig. 2D) [26], whose cleavage (C-PARP) is an apoptosis hallmark. Further confirmation of the lncRNA involvement in this axis was obtained in HD-MB03 cells using GapmeR #1, with *lncMB3* depletion leading to i) reduced RNA and protein levels for OTX2, NRL, CRX, and BCL2L1 (Fig. 3A and B); ii) decreased cell viability (Fig. 3C); and iii) corresponding modulation of C-PARP and PARP-1 levels (Fig. 3D).

**Figure 3.**
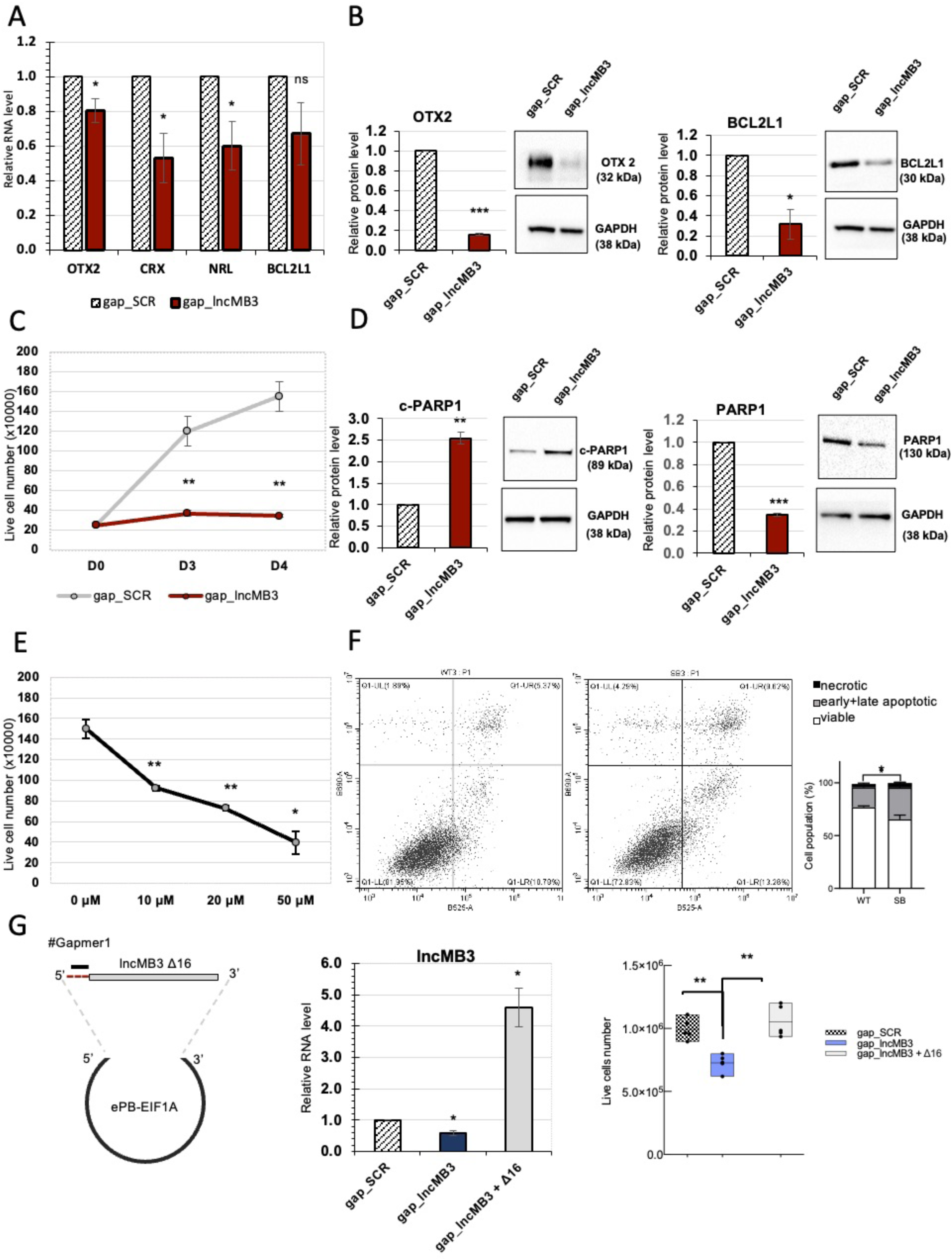
Analysis of TGF-β pathway and apoptosis in G3 MB cells. A. qRT-PCR analysis of *OTX2*, *NRL*, *CRX* and *BCL2L1* in HD-MB03 cells treated for 72 hours with gap_SCR or gap_*lncMB3* (GapmeR #1). Expression levels were compared to gap_SCR sample as control, set as 1. Data (means ± SEM) are expressed in arbitrary units and are relative to *GAPDH* mRNA levels. *N* = 3, * *p* ≤ 0,01 (two-tailed Student’s *t*-test). B. Western blot analysis of OTX2 (left panel) and BCL2L1 (right panel) protein levels in HD-MB03 cells treated for 72 hours with gap_SCR or gap_*lncMB3* (GapmeR #1). Normalisations were performed relative to GAPDH protein levels. N = 3, * *p* ≤ 0,05, *** *p* ≤ 0,001, (two-tailed Student’s t-test). C. Time course analysis of the number of viable HD-MB03 cells depleted for *lncMB3* (gap_*lncMB3*). Scramble-transfected cells (gap_SCR) were used as control. Cells were transfected at day 0 (D0) and counted at each timepoint. Data (means ± SEM) are expressed as the number of viable cells, counted by an automated cell counter. *N* = 4, ** *p* ≤ 0,01, (two-tailed Student’s *t*-test). D. Western blot analysis of c-PARP1 (left panel) and PARP1 (right panel) protein levels in HD-MB03 cells treated for 72 hours with gap_SCR or gap_*lncMB3* (GapmeR #1). *N* = 3, ** *p* ≤ 0,01, *** *p* ≤ 0,001. Details as in panel B. E. Dose-response analysis of the number of viable D283 Med cells after TGF-β inhibition. Control cells were treated with vehicle (DMSO). Cell counts started 24 hours after treatment. Data (means ± SEM) are expressed as the number of viable cells, counted by an automated cell counter. *N* = 4, * *p* ≤ 0,05, ** *p* ≤ 0,01 (two-tailed Student’s *t*-test). F. Annexin V staining of D283 Med cells upon TGF-β inhibition. Left panel: representative flow cytometry analysis of propidium iodide- and Annexin V-stained cells. Right panel: quantification of the fractions (expressed as percentage of total cell number) of viable (Annexin V-/PI-), early apoptotic (Annexin V+/PI-), late apoptotic (Annexin V+/PI+) and necrotic (Annexin V-/PI+) cells. For early+late apoptotic cells, *N* = 3, * p ≤ 0,05 (two-way ANOVA). **G.** Left panel: schematic representation of the construct overexpressing *lncMB3* Δ16. Middle panel: qRT-PCR analysis of *lncMB3* expression in D283 Med cells transfected with gap_SCR, gap_*lncMB3* or gap_*lncMB3* + Δ16 construct. *N* = 5. Details as in panel A. Right panel: number of viable D283 Med cells upon gap_*lncMB3* transfection (*lncMB3* KD) or Δ16 co-expression, compared to gap_SCR condition. Normalisation was performed on *lncMB3* KD condition. *N* = 5, ** *p* ≤ 0,01 (two-tailed Student’s *t*-test).

Consistent with MYC-regulation of *lncMB3* [10], RNA alterations seen with *lncMB3* KD were also observed in MYC-inhibited D283 Med cells *vs* untreated cells (Dataset 3). To further link *lncMB3* and the TGF-β pathway within the same regulatory network, we used a TGF-β Receptor Kinase inhibitor to block Smad2/3 phosphorylation. After 48 hour-treatment, we observed a dose-dependent reduction in viable cell number (Fig. 3E) correlated with increased apoptosis via Annexin V assay (Fig. 3F). These findings show that direct TGF-β cascade inhibition produced effects converging with *lncMB3* silencing.

To definitively confirm *lncMB3* role in counteracting apoptosis, we performed a rescue assay. A plasmid carrying the lncRNA sequence lacking its first 16 nucleotides (Δ16), necessary for GapmeR #1 recognition, was generated (Fig. 3G, left panel). When overexpressed in D283 Med cells along with the ASO, the mutant lncRNA (Fig. 3G, middle panel) restored cell viability compared to cells transfected with GapmeR #1 alone (Fig. 3G, right panel).

These findings reconstruct a molecular network initiated by MYC amplification and involving a non-coding RNA that suppresses apoptosis in G3 MB.

### Insights into the RNA interactome of *lncMB3*

RNA-RNA pairing is essential for lncRNA functionality. We investigated whether the lncRNA could influence TGF-β and apoptotic pathway genes through such interactions.

Initially, we hypothesized a straightforward interaction between *lncMB3* and candidate mRNAs downregulated upon its KD. However, preliminary analyses using the IntaRNA algorithm (http://rna.informatik.uni-freiburg.de/), which maps transcript binding regions, yielded low interaction propensity (Table 1). However, to experimentally explore indirect associations, we conducted RNA pull-down assays as in [14] (Fig. 4A). qRT-PCR analysis of *lncMB3*-enriched pull-down fractions with EVEN and ODD probe sets compared to LacZ-precipitated fractions (Table 2) showed no significant candidate enrichment (Table 3), excluding molecular interactions between *lncMB3* and TGF-β or downstream pathway transcripts.

**Figure 4.**
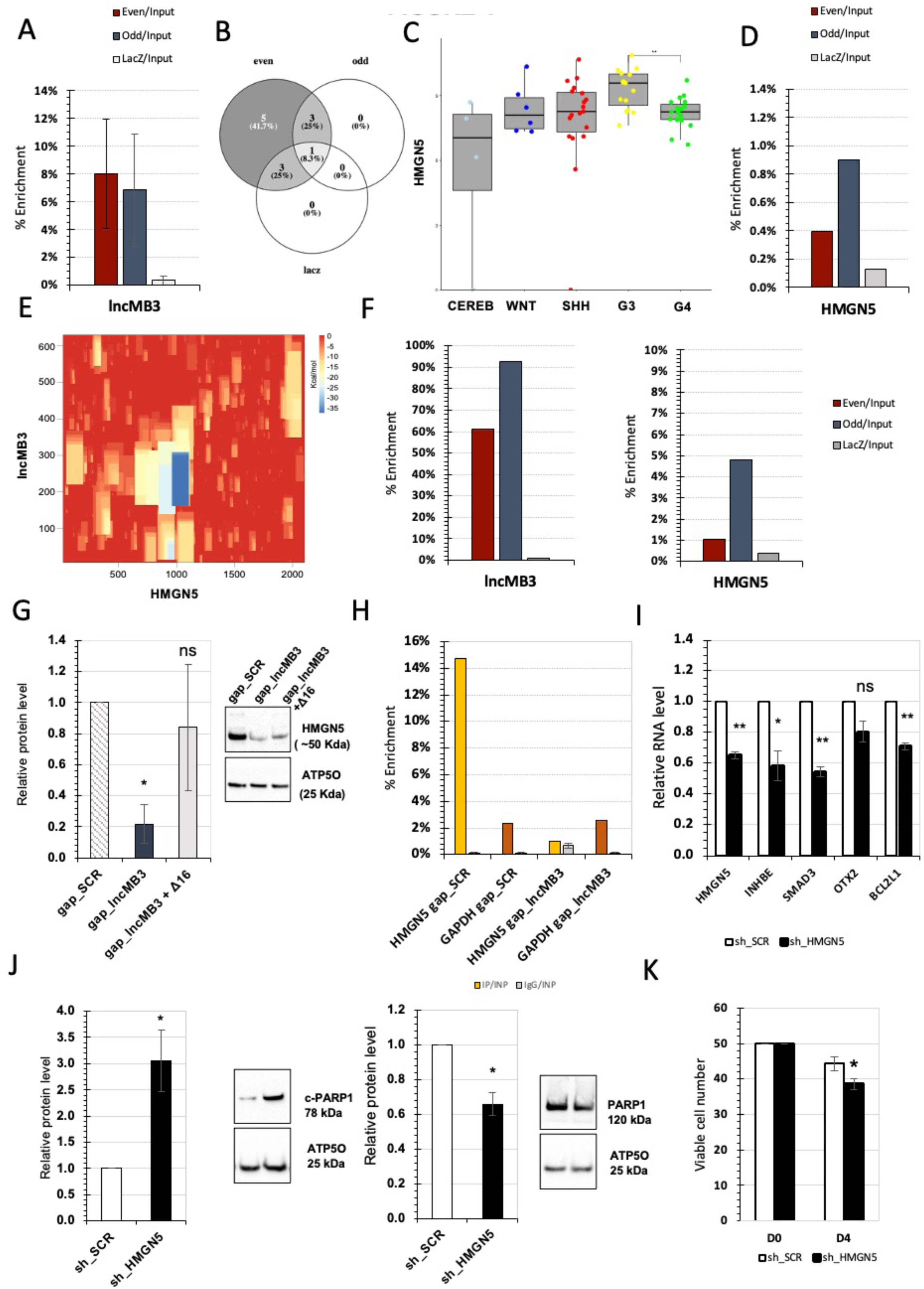
Analysis of *lncMB3*-*HMGN5* mRNA interaction. A. qRT-PCR analysis of RNA enrichment in EVEN, ODD and LacZ fractions over Input, from native *lncMB3* RNA pull-down experiments of D283 Med cell extracts. Data expressed as percentage of Input, *N* = 2. B. Venn diagram showing intersections of mRNAs retrieved and sequenced from two independent native *lncMB3* pull-down fractions (EVEN, ODD, LacZ). C. *HMGN5* mRNA expression in 4 healthy cerebella (normal samples) and 59 primary MB samples according to the transcriptomic dataset GSE164677. Results are expressed in log_2_(FPKM). * *p* ≤ 0,05, ** *p* ≤ 0,01, *** *p* ≤ 0,001, **** p ≤ 0,0001 (two-tailed Student’s *t*-test). D. qRT-PCR analysis of *HMGN5* mRNA enrichment in native *lncMB3* RNA pull-down fractions. Details as in panel A. E. IntaRNA energy map representing the predicted stability of RNA-RNA interaction between *lncMB3* and *HMGN5* mRNA. mRNA and lncRNA sequences are positioned along the x- and y-axes, respectively. The free energy of predicted intramolecular pairs ranges from red (higher energy, unstable pairing) to blue (minimal energy, stable pairing). F. qRT-PCR analysis of *lncMB3* (left panel) and *HMGN5* mRNA (right panel) enrichment in EVEN, ODD and LacZ fractions over Input, from AMT-crosslinked *lncMB3* RNA pull-down experiment in D283 Med cells. Data expressed as percentage of Input, *N* = 1. G. Western blot analysis of HMGN5 protein levels in D283 Med cells treated for 72 hours with gap_SCR or gap_*lncMB3* (GapmeR #1) or gap_*lncMB3* + Δ16 construct for *lncMB3* recovery. Normalisations were performed relative to GAPDH protein levels. N = 3, * *p* ≤ 0,05, (two-tailed Student’s t-test). H. qRT-PCR analysis of *HMGN5* and *GAPDH* upon RPL22-FLAG RIP assay in gap_SCR and gap_*lncMB3* (GapmeR #1) conditions. Analysis of RNA enrichment in IP and IgG fractions over Input. *N* = 1. Data expressed as percentage of Input. I. qRT-PCR analysis of *HMGN5*, *INHBE*, *SMAD3*, *OTX2*, *BCL2L1* mRNAs in D283 Med cells treated for 72 hours with sh_SCR or sh_HMGN5. Expression levels were compared to sh_SCR sample as control, set as 1. Data (means ± SEM) are expressed in arbitrary units and are relative to *GAPDH* mRNA levels. *N* = 3, * *p* ≤ 0,05, ** *p* ≤ 0,01 (two-tailed Student’s *t*-test). J. Western blot analysis of c-PARP1 (left panel) and PARP1 (right panel) protein levels in D283 Med cells treated for 72 hours with sh_SCR or sh_HMGN5. N = 3, * *p* ≤ 0,05 (two-tailed Student’s t-test). K. Analysis of number of viable D283 Med cells treated for 72 hours with sh_SCR or sh_HMGN5. Data (means ± SEM) are expressed as the number of viable cells, counted by an automated cell counter. *N* = 3, * *p* ≤ 0,05 (two-tailed Student’s *t*-test).

We then considered the competing endogenous RNA network, where long RNAs modulate microRNA activity by sponging them. Preliminary predictions of interactions between *lncMB3* and the core RNA-induced silencing complex component Argonaute 2 (AGO2), made using the *cat*RAPID algorithm (http://s.tartaglialab.com/page/catrapid_group) (Dataset 4) were experimentally validated through a CLIP assay (Fig. S5A). Based on these results, we employed Targetscan (https://www.targetscan.org/vert_80/) to identify microRNAs predicted to bind both *lncMB3* and candidate mRNAs (Table 4). Of six candidates (*hsa-miR-135b-5p*, *hsa-miR-216a-3p*, *hsa-miR-3681-3p*, *hsa-miR-613*, *hsa-miR-1197*, *hsa-miR-196b-5p*), the top three were tested by qRT-PCR on *lncMB3* pull-down fractions, showing no significant enrichment, ruling out competition for microRNA association (Table 5).

Alternatively, *lncMB3* may regulate target genes through interactions with other mRNAs [27]. We identified *lncMB3* interactome by cross-referencing RNA-Seq data from replicate independent *lncMB3* pull-downs (Fig. 4B and Dataset 5), applying a stringent analysis to pinpoint key binding partners. We found two mRNAs (*HMGN5*, *EIF5B*) and one lncRNA (*ANKDDA1*) associated with *lncMB3* (Fig. 4B and Fig. S6A). Notably, the mRNA for the nuclear factor HMGN5 (High Mobility Group Nucleosome Binding Domain 5) had the highest FC and the most significant FDR. Given HMGN5 broad epigenetic role in transcriptional regulation [28], its interplay with *lncMB3* highlights it a candidate for further analysis.

### *LncMB3* interacts with *HMGN5* mRNA

The chromatin factor HMGN5 regulates gene expression by influencing nucleosome-DNA interactions [28], [29]. *HMGN5* is ubiquitously expressed (Fig. S6B), with low expression in cerebellum, among brain tissues. Interestingly, RNA-Seq from the primary MB dataset GSE164677 showed *HMGN5* upregulation especially in G3 (Fig. 4C).

Co-enrichment of *HMGN5* mRNA with *lncMB3* was validated in an additional pull-down assay (Fig. 4D). IntaRNA predicted an interaction site (energy score: -30,1 kcal/mol) spanning ∼120 nucleotides between *lncMB3* 5’ region (nt 189-307) and *HMGN5* mRNA coding sequence (nt 933-1059) (Fig. 4E). This prediction was experimentally verified using a 4Z-aminomethyl-4,5Z,8-trimethylpsoralen (AMT)-crosslinked RNA pull-down [14], which detects direct RNA-RNA pairings *in vivo* (Fig. 4F). Interestingly, digital PCR-based absolute quantifications showed stoichiometrically equivalent expression levels of *lncMB3* and *HMGN5* (Fig. S6C).

To assess the functional consequences of this interaction, we investigated *lncMB3* influence on *HMGN5* expression. *HMGN5* mRNA levels remained unchanged following *lncMB3* KD, as per RNA-Seq data (Dataset 1) and qRT-PCR (Fig. S7A). However, HMGN5 protein levels dropped by ∼80% upon *lncMB3* KD (Fig. 4G), consistently with HMGN5 recognised oncogenic activity various cancers [30], [31]. Restoring *lncMB3* levels through transient Δ16 construct exogenous expression (Fig. 3G, left panel) showed a recovery trend for HMGN5 levels (Fig. 4G), indicating a specific and quantitative regulation of HMGN5 protein by *lncMB3*.

*LncMB3* impact on HMGN5 protein and their RNA interaction indicates it acts as a translational activator. This was confirmed using a Ribo-Tag strategy [15] with transient FLAG-tagged 60S ribosomal protein L22 (RPL22) expression in D283 Med cells (Fig. S7B). Western blot analysis of FLAG-RPL22-precipitated fractions showed comparable protein levels in gap_SCR *vs* gap_*lncMB3* extracts (Fig. S7C). At variance, qRT-PCR analysis revealed a marked decrease in *HMGN5* mRNA association with ribosomes in *lncMB3* absence, compared control (*GAPDH*, Fig. 4H). These results indicate that *lncMB3* is required for *HMGN5* mRNA ribosomal association.

### HMGN5 regulates TGF-β pathway and apoptosis in G3 MB cells

Given HMGN5 documented nuclear activity, we re-evaluated genes altered by *lncMB3* KD to assess if their expression changes were primarily transcriptional. qRT-PCR analysis was performed with exonic *vs* exon-intron junction primers to differentiate mature mRNAs from primary transcripts. As shown in Fig. S7D, *lncMB3* silencing downregulated *ACVR1B*, *INHBE*, *SMAD3*, *OTX2*, *NRL*, and *CRX* pre-mRNAs, highlighting a nuclear control over several genes regulated by *lncMB3* (compare Figs 1B, 1C, 2B, 3A).

HMGN5 role in suppressing apoptosis had not previously been assessed in MB. To functionally connect *lncMB3* and HMGN5 activities, we aimed to mimic the molecular effects of *lncMB3* KD by silencing *HMGN5*. Using VectorBuilder (https://en.vectorbuilder.com/) we designed a short interfering RNA targeting *HMGN5*. Transient transfections in D283 Med cells resulted in a ∼30% downregulation of *HMGN5* mRNA (Fig. 4I) and a ∼60% reduction in protein levels (Fig. S7E). This led to a 20-50% RNA decrease for *INHBE*, *SMAD3*, *OTX2*, and *BCL2L1* (Fig. 4I), targets also downregulated upon *lncMB3* KD (see Figs 1B, 1C, 2B, 3A), indicating that *HMGN5* KD appreciably phenocopy *lncMB3* molecular effects (Fig. S7F). Regarding cell death, *HMGN5* KD caused a threefold increase in c-PARP1 protein (Fig. 4J, left panel) and a 30% decrease in full-length PARP1 (Fig. 4J, right panel). HMGN5 level reduction also mirrored a viable D283 Med cell number decrease, with a drop of ∼30% (Fig. 4K), similar to the impact of *lncMB3* targeting. These findings demonstrate that *lncMB3* and HMGN5 participate into the same molecular and cellular phenotypes. Dependent on interactions with *lncMB3*, *HMGN5* mRNA decreased translation contributes to the nuclear regulation of TGF-β pathway and downstream gene module in G3 MB cells.

### *LncMB3* inhibition synergizes with DDP treatment in G3 MB cells

Cisplatin (DDP) is a first-line anti-cancer agent and a cornerstone of MB treatment [32]. Combining therapies can reduce DDP side effects and fight both intrinsic and acquired resistance. Given the shared influence of *lncMB3* and DDP on apoptosis-related pathways [33], we asked whether *lncMB3* KD could amplify DDP cytotoxicity in G3 MB cells.

Initial tests of DDP administration to D283 Med cells (1-10μM) reduced viable cell number by 10%-20% after 48 hours (Fig. 5A, left panel). We then combined 10μM DDP with 100nM GapmeR #1 or GapmeR #2 transfection and counted surviving cells. Normalising double-treated *vs* drug-untreated sample cell counts revealed lower relative survival rate in gap_*lncMB3*-treated cells compared to scramble-transfected, indicating a synergistic effect between treatments (Fig. 5A, right panel). This was confirmed through an apoptosis assay with Annexin V staining (Fig. 5B, left panel). Quantification of non-viable *vs* viable cells showed an increase of late apoptotic cell number compared to other populations upon combined treatments (Fig. 5B, middle panel), reinforcing the synergistic effect (Fig. 5B, right panel).

**Figure 5.**
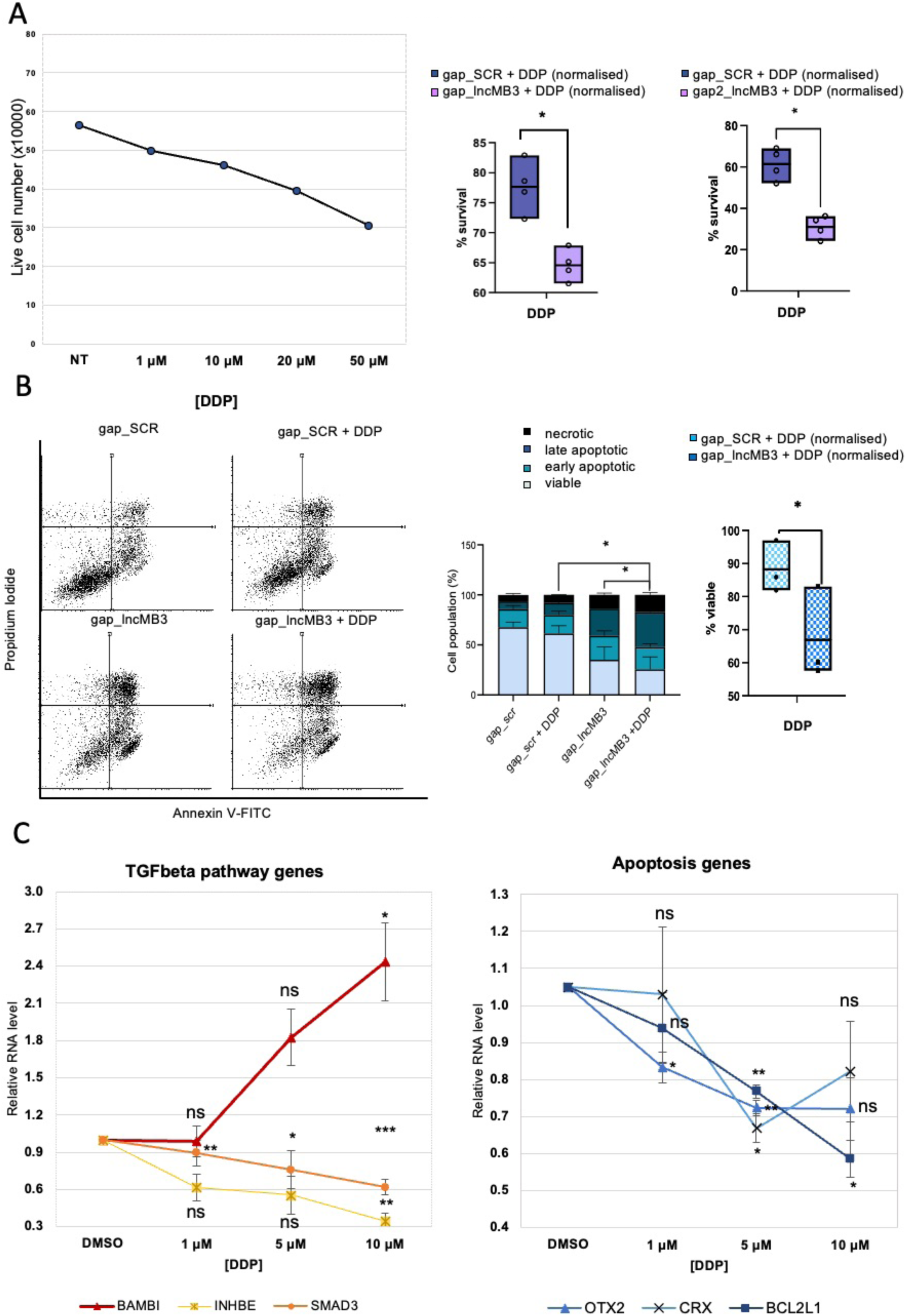
Analysis of *lncMB3* targeting and DDP treatment on D283 Med cell viability and apoptosis. A. Left Panel: Dose-response analysis of the number of viable D283 Med cells upon DDP treatment. Control cells (NT) were treated with the DDP vehicle (DMSO). Cell counts were conducted 24 hours after DDP administration. *N* = 1. Middle Panel: percentage of viable D283 Med cells following DDP administration and *lncMB3* GapmeR transfection (GapmeR #1, middle panel; GapmeR #2, right panel). Cell counts were conducted 48 hours after DDP/GapmeR treatments. Double-treated samples (gap_SCR+DDP and gap_*lncMB3*+DDP) were normalised on the viable cell number of the corresponding transfected-only sample (gap_scr and gap_*lncMB3*, respectively). DDP concentration used was 10μM. *N* = 4, * p ≤ 0,05 (two-tailed Student’s t-test). B. Apoptosis analysis of D283 Med cells following DDP administration [1μM] and GapmeR #1 transfection. Left panel: representative flow cytometry analysis of PI- and Annexin V-stained D283 Med cells 24 hours after *lncMB3* KD, with or without DPP treatment. Middle panel: quantification of viable (Annexin V-/PI-), early apoptotic (Annexin V+/PI-), late apoptotic (Annexin V+/PI+) and necrotic (Annexin V-/PI+) fractions. *N* = 3, * p ≤ 0,05 (comparison between early + late apoptotic cells). Right panel: percentage of viable D283 Med cells. Data normalisations as in (A). *N* = 3, * p ≤ 0,05 (two-way ANOVA). C. qRT-PCR analysis of TGF-β pathway genes (left panel) and downstream pathway genes (right panel) in D283 Med cells treated for 24 hours with DDP at different doses (reported on the x-axis). Control cells were treated with vehicle (DMSO) and set as 1. Data (means ± SEM) are expressed in arbitrary units and are relative to *GAPDH* mRNA levels. *N* = 3, * *p* ≤ 0,05, ** *p* ≤ 0,01, *** *p* ≤ 0,001 (two-tailed Student’s *t*-test).

To examine whether synergy relied on a crosstalk between treatments we tested DDP administration consequence on *lncMB3* target genes. A partial deregulation of both the TGF-β (Fig. 5C, left panel) and apoptotic pathway gene (Fig. 5C, right panel) was highlighted, similar to *lncMB3* KD. *LncMB3* expression remained unaffected by chemotherapy (Fig. S8A), suggesting the two treatments as independent triggers with overlapping outcomes.

The synergistic enhancement of anticancer effects was not indiscriminate. When GapmeR #1 was combined with vincristine, another MB therapeutic drug [34], only additive, not synergistic effects were measured on viable cell counts (Fig. S8B) and apoptosis (Fig. S8C). These data suggest that, *in vitro*, *lncMB3* targeting potentiates DDP treatment effectiveness in reducing G3 MB cell proliferation and inducing apoptosis.

### Production of the ferritin variant HFt-HIS-PASE

Targeting *lncMB3* affects the TGF-β pathway, enhances apoptosis, and boosts chemotherapeutic efficiency in cancer cell lines. This led us to explore the use of human ferritin nanoparticles (NPs) to formulate innovative RNA/protein complexes aimed at *lncMB3*-directed GapmeR delivery. Human ferritin H-type (HFt) is a promising NP for drug delivery, particulary chemotherapeutics, due to its internalisation by the transferrin receptor 1 (CD71), frequently overexpressed in tumours [13].

We designed and synthesized HFt-HIS-PASE, a recombinant ferritin variant derived from a prior HFt version, HFt-MP-PASE [35]. HFt-HIS-PASE contains a tumour-selective sequence (MP), responsive to tumour protease MMP 2 and 9 activity, and a shielding polypeptide (PASE) that enhances stability, masks the surface, and increases target specificity. Additionally, HFt-HIS-PASE features a five-histidine motif known to aid nucleic acid endosomal escape of [36] (Fig. S9A).

Preliminary qRT-PCR assessments of *MMP 2*, *MMP 9*, and *CD71* mRNA abundance in HD-MB03 and D283 Med cells (Fig. S9B) led to selecting the latter for further tests. We then evaluated the targeting capacity of a Fluorescein-5-Maleimide-labeled HFt-HIS-PASE nanovector (HFt-HIS-PASE-fluo). About 85% of D283 Med cells exposed to HFt-HIS-PASE-fluo (1mg/mL) showed fluorescence signal 12 hours post-administration, indicating HFt-HIS-PASE affinity (Fig. 6A). Immunofluorescence assays performed 24 hours later, using nuclear (DAPI) and cytoplasmic (Phalloidin) staining and confocal microscopy analysis, demonstrated intracellular uptake of HFt with a nuclear-cytoplasmic distribution (Fig. 6B).

**Figure 6.**
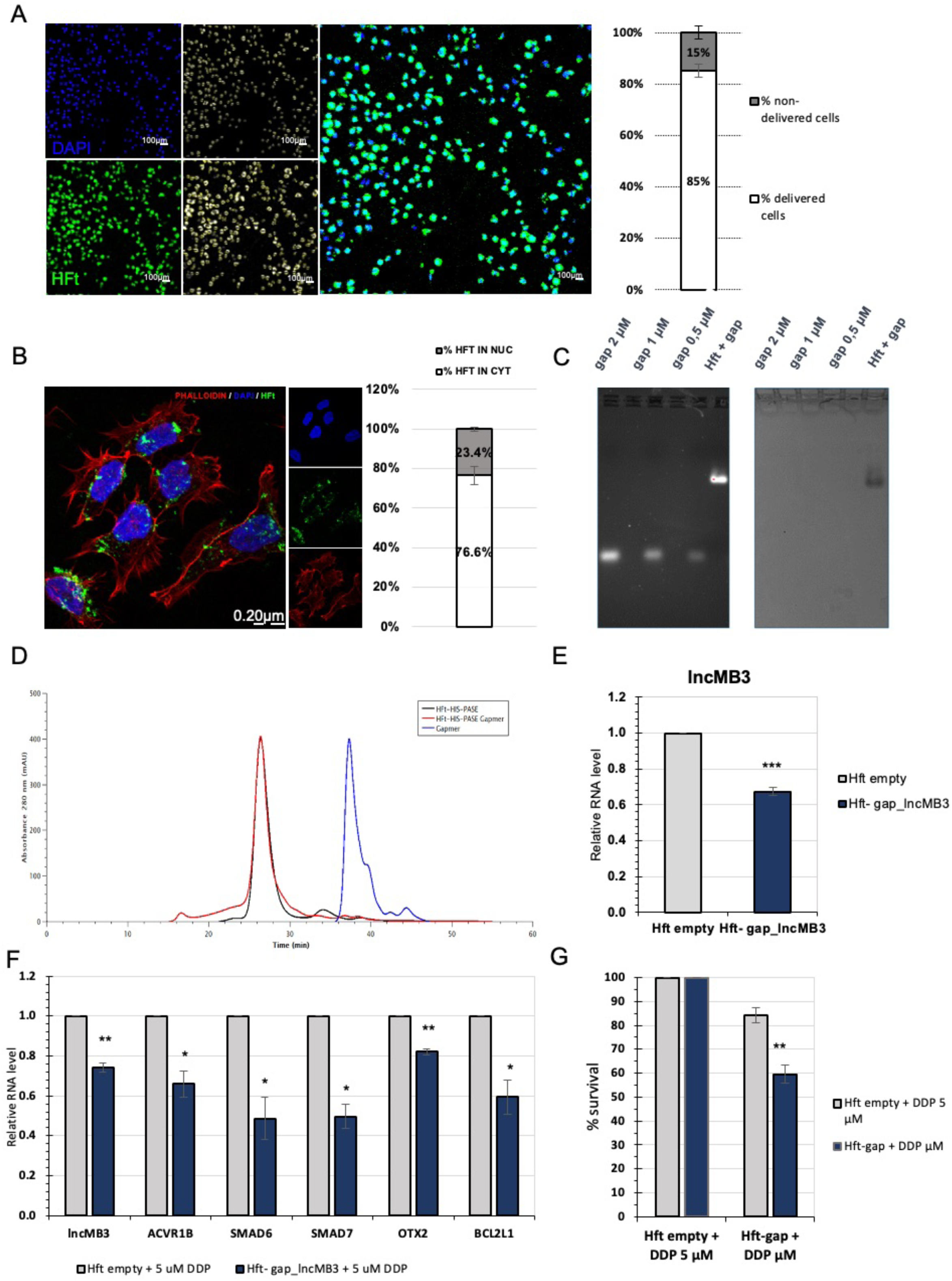
Synthesis, analysis and application of HFt-HIS-PASE/GapmeR complexes. A. Left panel: representative image of colocalisation analysis between fluorescein-HFt-HIS-PASE and DAPI in D283 Med cells, 12 hours after administration (magnification 20x). Fluorescein-HFt-HIS PASE (green) and DAPI (blue, nuclear staining) are on the left, object counts after binarisation are in the middle and the merge of the two channels is on the right. Right panel: percentage of colocalisation between fluorescein-HFt-HIS-PASE and DAPI in D283 Med cells. Data (mean ± SEM) were obtained from 8 different fields. B. Left panel: representative immunofluorescence of fluorescein-HFt-HIS PASE delivered for 24 hours (green), DAPI (blue, nuclear staining) and phalloidin (red, cytoplasmic staining) (magnification 60x). Right panel: percentage of nucleus/cytoplasm subcellular localisation of HFt-HIS PASE in D283 Med cells. Data (mean + SEM) were obtained from 5 different fields. C. Band migration profiles on agarose gel electrophoresis. Gel was double stained with SYBR Gold for GapmeR visualisation (left panel) and Coomassie Blue for ferritin visualisation (right panel). *Lane 1*: GapmeR standard 2µM; *Lane 3*: GapmeR standard 1µM; *Lane 5*: GapmeR standard 0,5µM; *Lane 6*: HFt-HIS-PASE-GapmeR complex. D. Size Exclusion Chromatography profile analysis of HFt-HIS-PASE (in black), HFt-HIS-PASE-GapmeR #1 complexes (in red) and GapmeR #1 alone (in blue). E. qRT-PCR analysis of *lncMB3* levels in D283 Med cells upon treatment with HFt-HIS-PASE-GapmeR #1 complexes, 48 hours after delivery, compared with empty HFt-HIS-PASE, set as 1. Data (means ± SEM) are expressed in arbitrary units and are relative to *GAPDH* mRNA levels. *N* = 3, *** *p* ≤ 0,001 (two-tailed Student’s *t*-test). F. qRT-PCR analysis of TGF-β pathway genes in D283 Med cells treated for 48 hours with HFt-HIS-PASE-GapmeR #1 + 5µM DDP. Expression levels were compared to HFt-HIS-PASE-empty + 5µM DDP, set as 1. Data (means ± SEM) are expressed in arbitrary units and are relative to *GAPDH* mRNA levels. *N* = 3, * *p* ≤ 0,05, ** *p* ≤ 0,01 (two-tailed Student’s *t*-test). G. Analysis of number of viable D283 Med cells treated for 48 hours with HFt-HIS-PASE-GapmeR #1 + 5µM DDP. HFt-HIS-PASE empty + 5µM DDP was used as control condition. Data (means ± SEM) are expressed as the percentage of viable cells, counted by an automated cell counter. *N* = 4, ** *p* ≤ 0,01 (two-tailed Student’s *t*-test). Where necessary in the figure HFt-HIS-PASE is referred as HFt.

### Analysis of HFt-HIS-PASE-GapmeR complexes and *lncMB3* targeting

To evaluate *lncMB3* targeting using HFt-ASO complexes, we generated and characterised HFt-HIS-PASE-GapmeR nucleoprotein particles. The *lncMB3* GapmeR #1 was encapsulated in the internal cavity of HFt-HIS-PASE via pH-dependent HFt dissociation/reassociation. Successful GapmeR loading was confirmed by electrophoresis of purified HFt-HIS-PASE-GapmeR complexes. Distinct migration patterns for the HFt-HIS-PASE-GapmeR complex compared to free GapmeR were observed. DNA-specific staining (Fig. 6C, left panel) displayed significant delay for nucleic acid when associated with HFt-HIS-PASE (lane HFt + gap), while protein-specific staining confirmed a co-migration profile between HFt-HIS-PASE-GapmeR complex and ferritin (Fig. 6C, right panel).

Regarding stability, no nucleic acid degradation was observed after 1 hour and overnight incubations of HFt-HIS-PASE-GapmeR with DNA/RNA nuclease at 37°C (Fig. S9C). No degradation was noted after 3 months of storage at 4°C, indicating GapmeR encapsulation within the protein cavity, shielded from external exposure.

To evaluate GapmeR binding efficiency to HFt NPs, of agarose band relative intensities of DNA or protein molecules were quantified. Comparisons were made with free GapmeR and HFt-HIS-PASE molecules at standard concentrations. The final GapmeR-HFt-HIS-PASE molecular ratio was 0,5:1. Purity and hydrodynamic volume of HFt-HIS-PASE-GapmeR were determined by size-exclusion chromatography, showing similar elution volume and size to non-encapsulated HFt-HIS-PASE protein, with no free GapmeR detected, indicating co-elution with ferritin (Fig. 6D). The biological activity of the complex was tested in D283 Med cells treated with a single dose of HFt-HIS-PASE alone or GapmeR-complexed (200nM). Steady-state *lncMB3* levels, 48 hours post-treatment, showed a 30% reduction with encapsulated GapmeR #1 compared to mock-treated cells (Fig. 6E). Similar outcomes were obtained with HFt-HIS-PASE-GapmeR stored at 4°C for 3 months, replicating the effects of a standard, single-pulse GapmeR #1 transfection at 100nM for 48 hours (Fig. S9D). No modulation of TGF-β pathway gene expression, cell proliferation reduction, or apoptosis increase was registered under these conditions, probably due to insufficient *lncMB3* downregulation. However, to further investigate HFt-HIS-PASE-mediated cellular effects of *lncMB3* targeting, we leveraged our findings on the synergy between *lncMB3* KD and DDP administration. D283 Med cells were treated with 5μM DDP alone or in parallel to HFt-HIS-PASE-GapmeR. Combined treatments led to a 20-50% reduction in TGF-β pathway and downstream genes, (Fig. 6F), as revealed by qRT-PCR analysis. A decrease in cancer cell survival was also observed, compared to DDP treatment alone (Fig. 6G), consistent with earlier data (Fig. 5A), indicating that HFt-based NPs are effective for *lncMB3* targeting and sensitising G3 MB cells to anticancer drugs.

## DISCUSSION

Integrated multi-omics has combined molecular genetic with histological analyses within a framework to illuminate MB inherent heterogeneity and complexity [1]. Despite unique genomic, biochemical and clinical traits in each MB subgroup [37], therapies still rely on general interventions -surgical resection, chemotherapy, and radiotherapy-with 5-year survival rates of 60%-80% [38]. This underscores the need for targeted therapies and deeper explorations of dysfunctional pathways, particularly in high-risk and poorly characterised subgroups, like G3 and G4. G3 MB, representing 25% of cases, exhibits the highest recurrence and metastasis rate, correlating with worse outcomes [39]. The discovery of MYC amplification/overexpression as a major pathogenic factor [40], [41] aligns with estimates of aberrant MYC activity in 70% of human cancers [41]. Yet, targeting MYC has long been challenging due to its disordered structure, leading to low-potency therapeutic candidates, with suboptimal pharmacological properties and considerable off-target effects [42]. Furthermore, MYC critical role in physiological processes which are central to development and post-natal life (differentiation, proliferation, and cell cycle regulation), its inhibition in paediatric cancers must be tackled. Several pre-clinical studies and clinical trials are investigating direct MYC inhibitors [42], [43], like OMOMYC, a dominant-negative showing promise in solid cancers [10], [43], that we used to map the MYC transcriptome in G3 MB cells [10]. Indirect strategies targeting MYC cofactors or regulators are valuable alternatives and, along this direction, understanding MYC-dysregulated gene pathways, including non-coding RNA networks, may diversify and improve opportunities for MYC-targeted therapies [44].

Over the last two decades, the rise of non-coding RNAs has reshaped cancer biology [45]. LncRNAs have gained attention as cancer hallmark modulators for their tissue- and cell-specific expression and structural flexibility, making them attractive biomarkers and molecular targets. However, their potential as biological master regulators require detailed functional characterisation.

To date, only few lncRNAs have been mechanistically described in G3 MB. This study clarifies the role of MYC-regulated *lncMB3* in G3 medulloblastomagenesis, establishing a foundation for its potential application. *LncMB3* is upregulated in G3 MB cell lines and primary tumours, playing a pivotal role in cell death evasion *in vitro*. Through transcriptomic, molecular, and cellular assays, we show that *lncMB3* regulates the TGF-β cascade, crucial for G3 MB pathology, impacting cancer cell proliferation and survival [11]. Notably, we identified *lncMB3* target gene cluster in this pathway, whose mysregulation elevates the retinal TF OTX2, an oncogenic driver in G3 MB that boosts cell proliferation, suppresses apoptosis, and contributes to tumour aggressiveness [46]. OTX2 and other photoreceptor TFs, including NRL and CRX, drives anti-apoptotic factors like BCL2L1 [25], linking this gene program to sustained cancer cell viability. *LncMB3* activity orchestrates key pathways, genes, and processes in G3 MB, forming a nexus among *MYC* driver gene, TGF-β signalling, photoreceptor gene network, and apoptosis regulation. This interconnection offers an integrated framework for understanding the molecular underpinnings of this tumour subgroup.

While our coding transcriptome analysis provided insights into *lncMB3* function, its interactome revealed its mode of action. *LncMB3* directly interacts with HMGN5 mRNA, encoding an epigenetic regulator within the HMGN nucleosome-binding and nuclear architecture family, involved in extensive chromatin decondensation and transcriptional regulation [30], [31]. *HMGN5*, implicated in embryonic gene regulation [47] and exhibiting oncogenic properties in several cancers [30], [31], is upregulated in G3 MB. Our findings suggest that functional RNA-RNA interactions between coding and non-coding molecules -an emerging theme in cancer regulation-upregulate HMGN5 protein levels, explaining why *HMGN5* RNA interference phenocopies *lncMB3* targeting effects. Given that several *lncMB3* targets are dysregulated in the nucleus, where HMGN5 operates, this mechanism adequately accounts for the observed molecular and cellular phenotypes, though additional regulative cascades or secondary effects could exist, consistent with multifunctional scaffolding properties and diverse subcellular localization of lncRNAs. Finally, *lncMB 3*/*HMGN5* co-expression confined to pathological cerebellum suggests their interaction as a platform for developing novel inhibitors competing out disease-specific complexes.

Due to G3 MB aggressive nature and treatment resistance, integrated therapies promise solutions. ASOs have gained prominence in RNA-based treatments for their design flexibility and pharmacological properties [45]. Our analysis of *lncMB3*-targeting LNA GapmeRs combined with standard MB chemotherapy demonstrated synergistic interplay enhancing drug cytotoxicity. This effect likely stems from both treatments acting on the intrinsic apoptotic pathway [33] and the TGF-β cascade (this study). These findings suggest modulating the *lncMB3* molecular network to amplify anticancer agent efficacy, supporting RNA-based combinatorial strategies to reduce tumour resistance and improve G3 MB treatment outcomes. However, a major challenge in RNA therapies is ensuring efficient effector delivery to intended sites. We explored application of HFt-based nanocarriers for deploying LNA GapmeRs targeting *lncMB3*. HFt properties, from high biocompatibility and stability to low toxicity and cost-effectiveness, make it a powerful drug delivery system [13]. Moreover, HFt capacity to cross the blood-brain barrier [13] positions it as a promising tool for brain tumour therapies. Its symmetrical self-assembly, small and uniform size, and versatile surface functionalisation are ideal for bioactive compound delivery, including small nucleic acids. While siRNAs [48] and microRNAs [49] have already been delivered by HFt, this studu is the first, to our knowledge, to employ HFt-encapsulated ASOs for gene silencing. Combinatorial administration of *lncMB3*-targeting NPs and cisplatin replicated the synergistic effects observed with ASO transfection, impacting the TGF-β pathway and reducing cell survival. These results indicate that HFt-based strategies have potential for targeting oncogenic pathways at the RNA level via biocompatible delivery of tumour-specific antisense oligomers.

In conclusion, this study highlights the critical role of *lncMB3* in G3 MB apoptosis regulation through TGF-β pathway modulation and interaction with underexplored genes such as HMGN5. *LncMB3* stands out as a key regulatory node linking MYC amplification to enhanced tumour cell survival via cell death inhibition (Fig. 7). Future efforts will focus on deeper exploration of *lncMB3* interactions and regulatory mechanisms-of-action, as well as validating its therapeutic potential in preclinical G3 MB models using the HFt nanocarriers.

**Figure 7.**
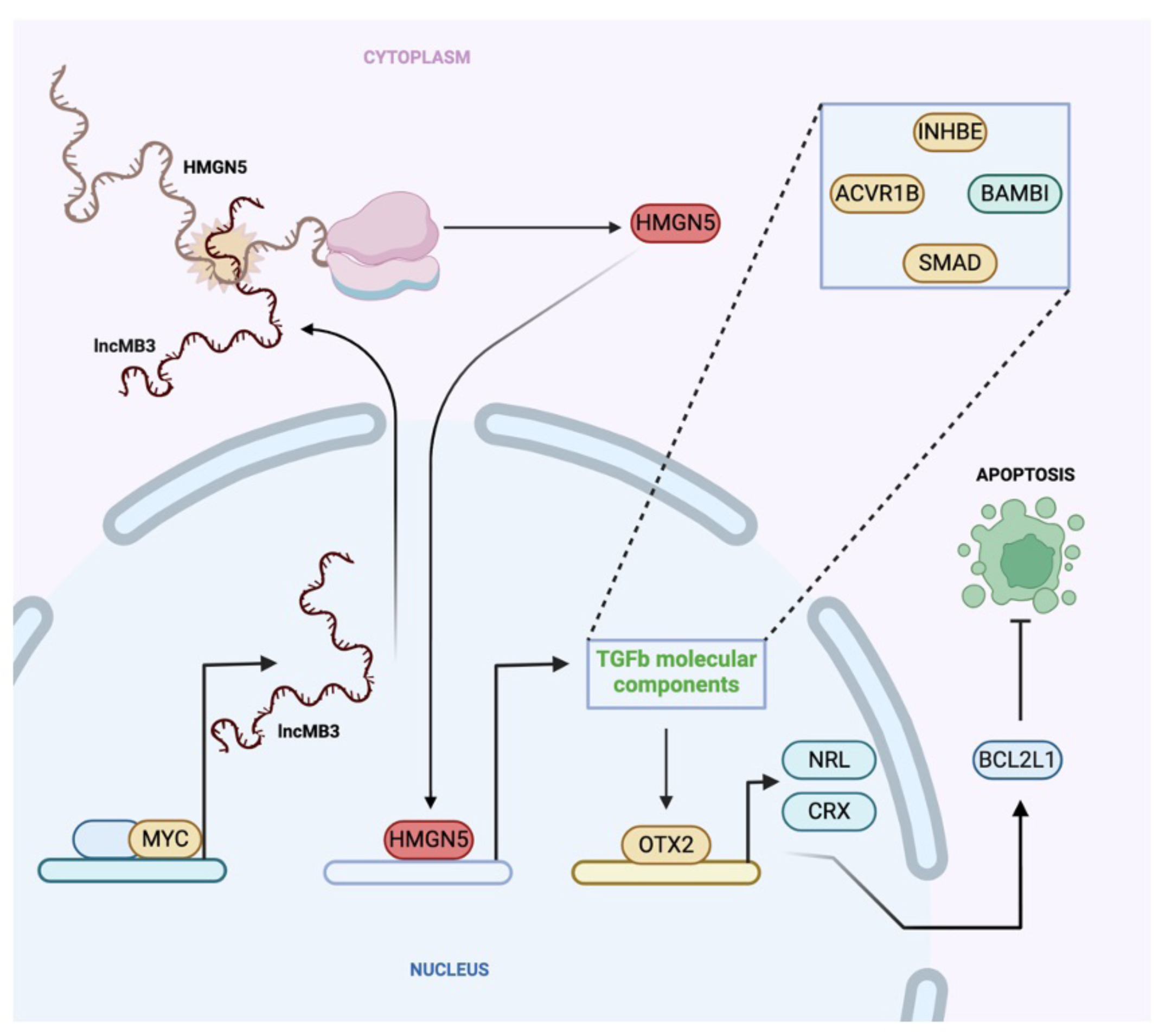
Regulatory circuit. Schematic representation of the MYC-dependent *lncMB3* mechanism of action. Image made with Biorender (https://biorender.com)

## ACKNOWLEDGEMENTS

We acknowledge M. Marchioni and J. Genovese for their experimental assistance, C. Grelloni and A. Giuliani for their technical and analytical suggestions and E. Caffarelli for discussions and laboratory support. FRP was supported by an AIRC fellowship. Confocal microscopy acquisitions were conducted at the Imaging Facility of CLN²S@Sapienza, Istituto Italiano di Tecnologia, Rome. Cytofluorimeters were available at IBPM-CNR. RNA-Seq were performed by DNA Link (Seoul, KR) and by Eclipse Bioinnovation (San Diego, CA).

In memory of M. Arceci, beloved friend and colleague.

## AUTHOR CONTRIBUTIONS

CRedIT author statement. Conceptualisation: AG, PL; Investigation: AG (gene expression analysis, cloning, molecular and cellular assays), PT (interaction assays, imaging and microscopy), FRP (flow cytometry), EF and GT (nanovector synthesis and characterisation), AB (gene expression analysis), AP, FM and AC (bioinformatic analysis); Formal Analysis: AG, PT, FRP, AP, FM, AB and AC; Validation: AG, PT, FRP, EF, AB and GT; Methodology: AG, PT, FRP, EF, JR and GT; Data Curation: AG, PT, AP, FM and AC; Software; AP, FM and AC; Visualisation: AG and JR; Resources: AC, IB, DT, MB and EF; Supervision: AC, DT, MB, PC and PL; Writing – original draft (main text): AG, MB and PL; Funding Acquisition and Project administration: PL.

## DATA AVAILABILITY

The RNA-Seq data presented in this study are available in GEO, accession numbers GSE277976 (token: iredyqecjjsvzqp) and GSE278666 (token: gdmfuqsghbmnxoj).

## INFORMED CONSENT

Manuscript is approved by all authors for publication.

