## Supplementary figures and images for "A circuit involving the lncRNA MB3 and the driver genes MYC and OTX2 inhibits apoptosis in Group 3 Medulloblastoma by regulating the TGF-β pathway via HMGN5"

### SUPPLEMENTARY FIGURE 1

#### FIGURE S1

## S1A

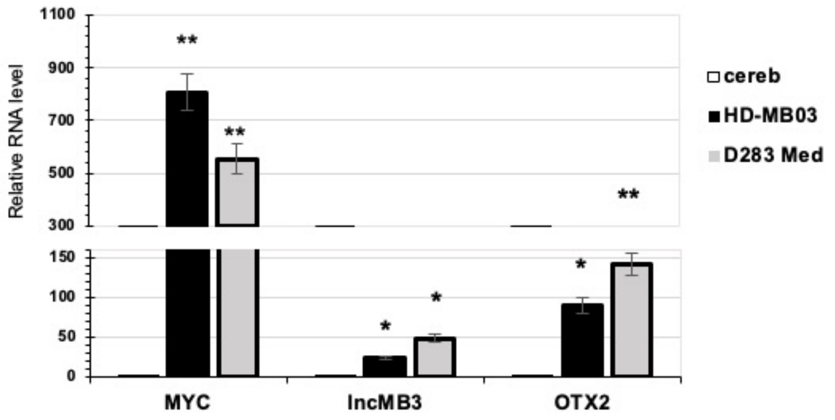


## S1B

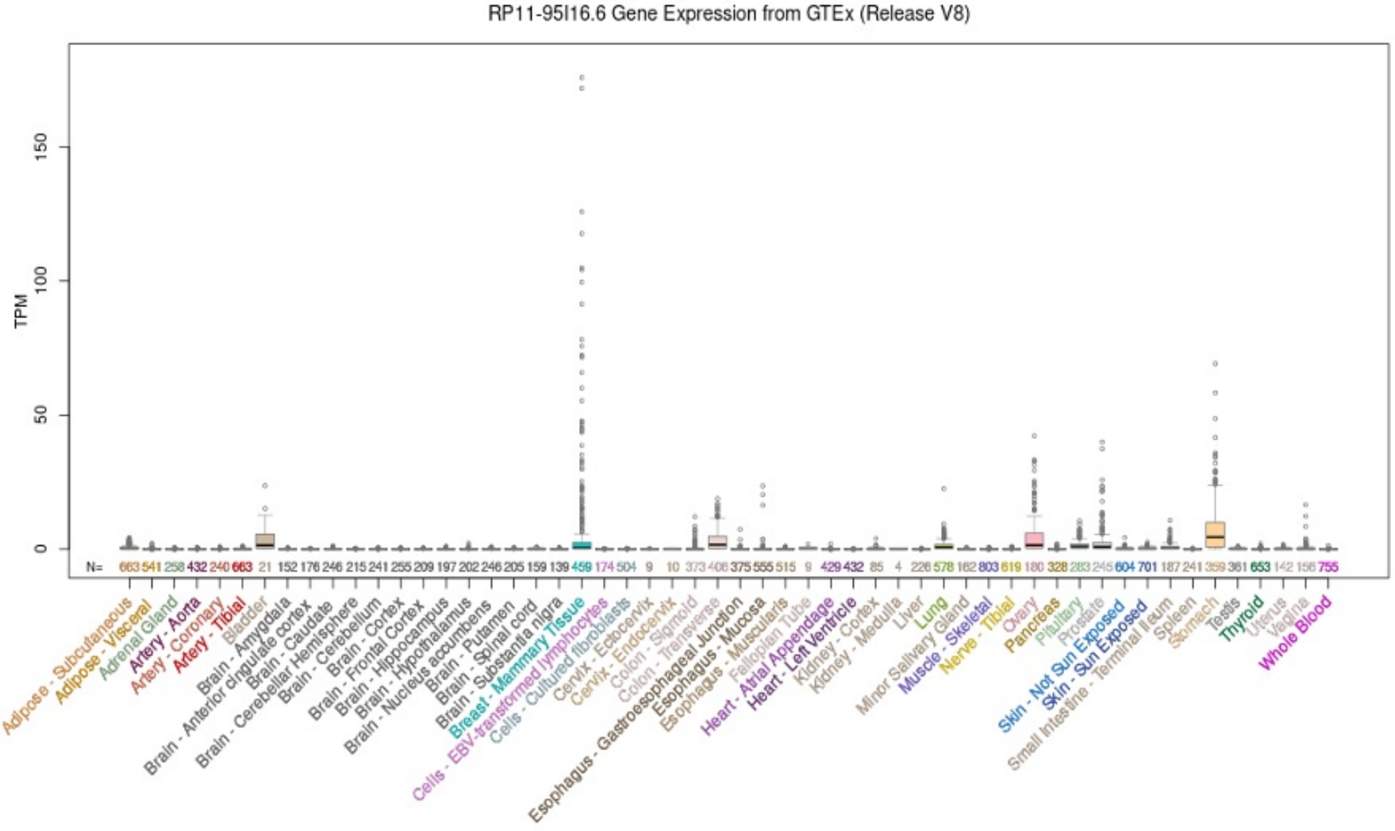

### SUPPLEMENTARY FIGURE 2

#### FIGURE S2

S2A

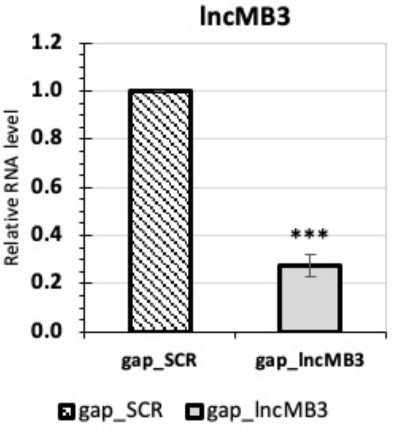


S2B

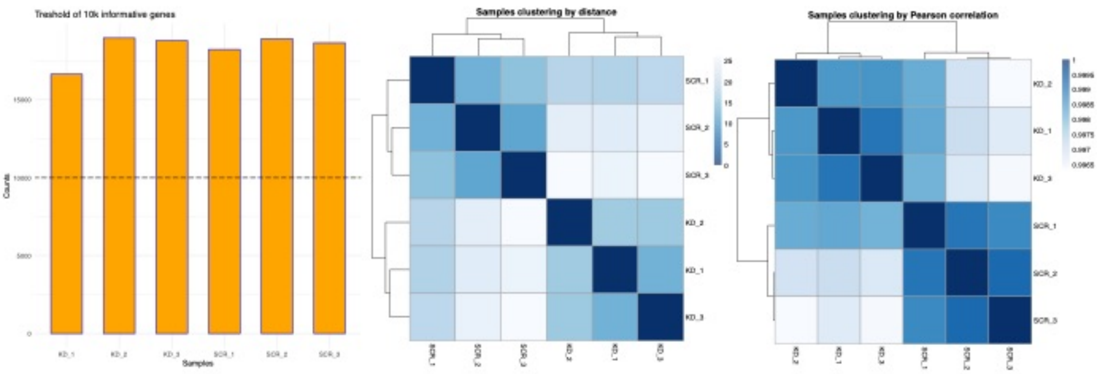


S2C

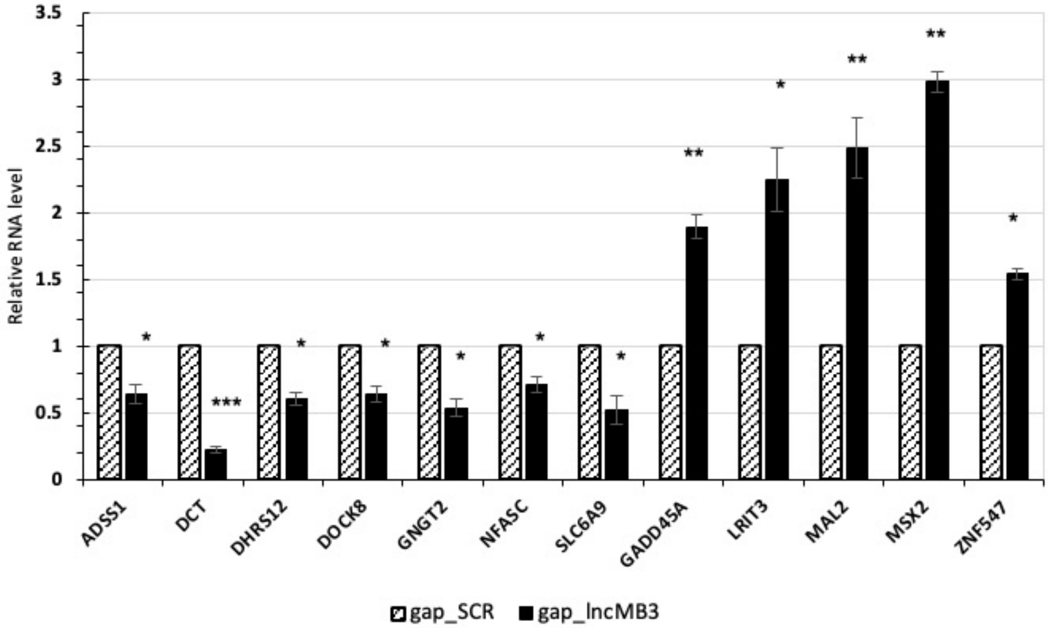


S2D

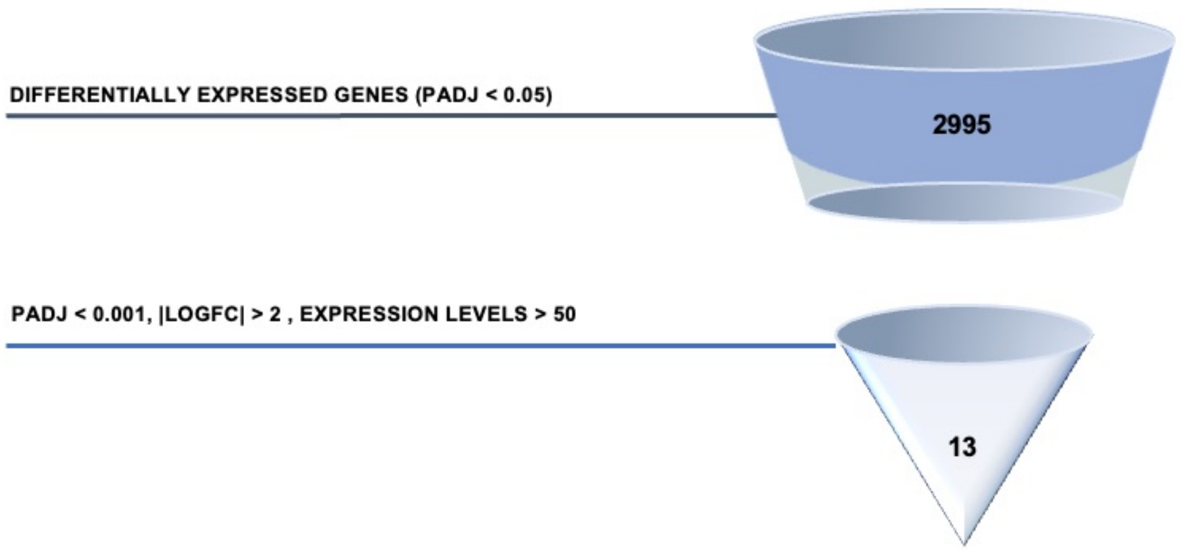

### SUPPLEMENTARY FIGURE 3

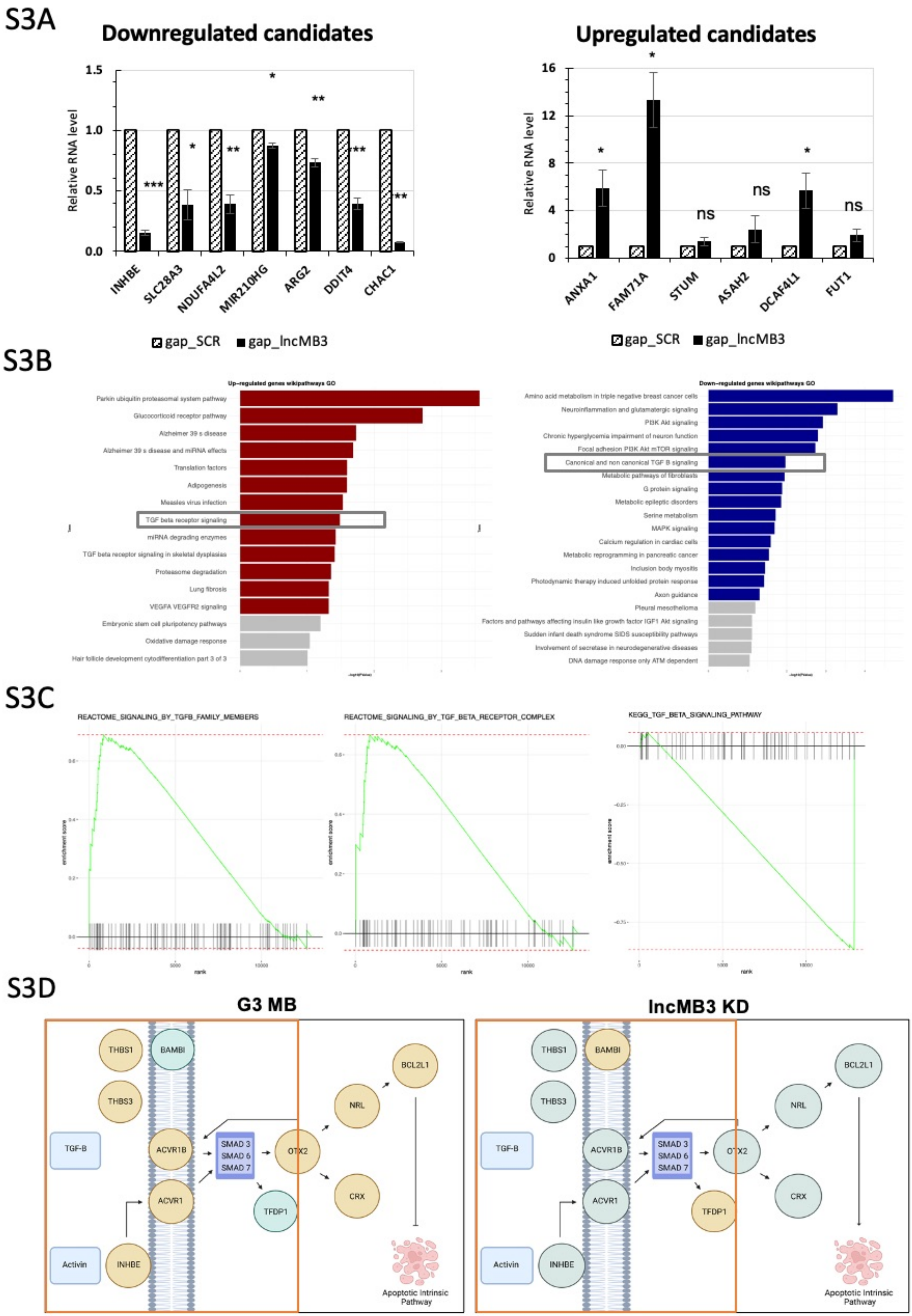

### SUPPLEMENTARY FIGURE 4

#### FIGURE S4

## S4A

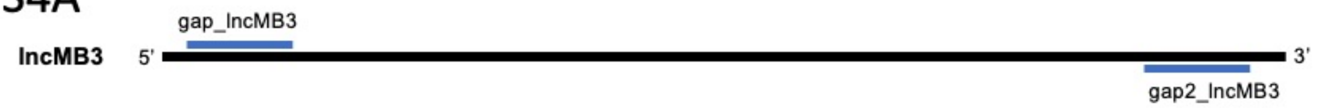


## S4B

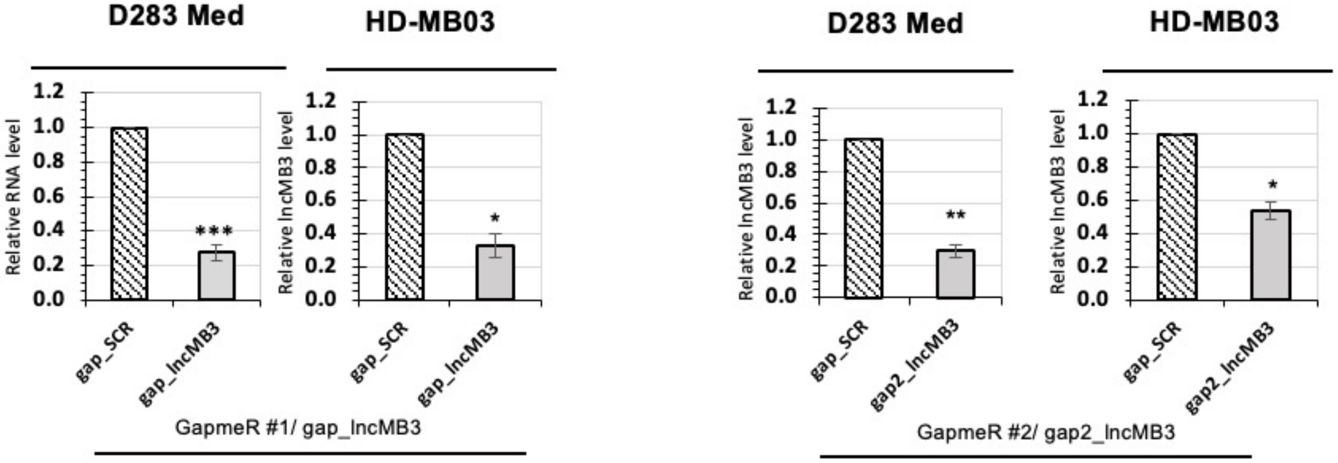

### SUPPLEMENTARY FIGURE 5

#### FIGURE S5

## S5A

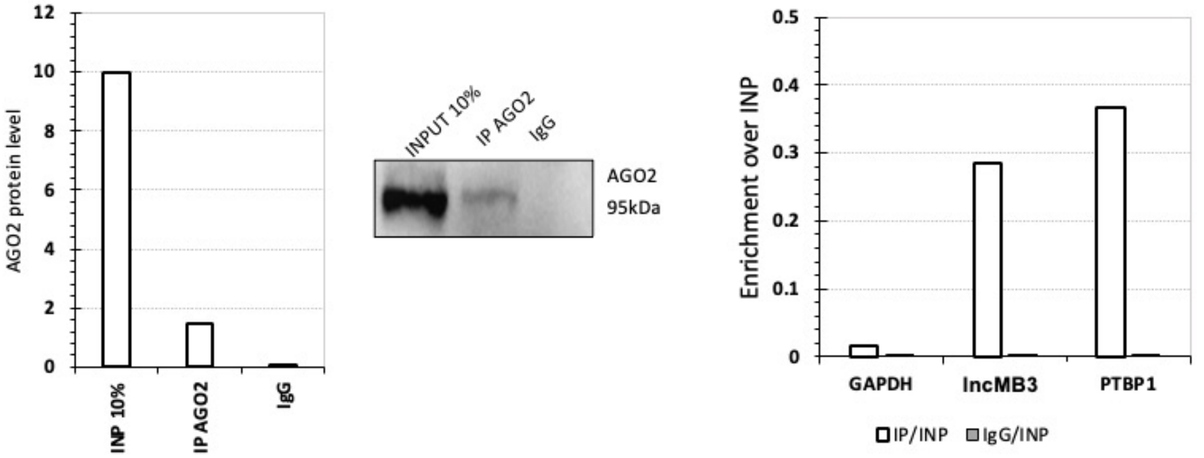

### SUPPLEMENTARY FIGURE 8

#### FIGURE S8

## S8A

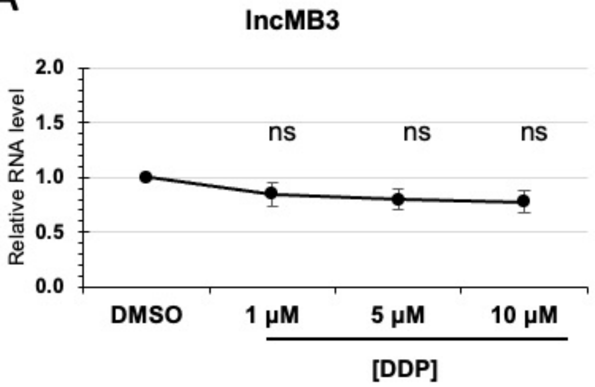


## S8B

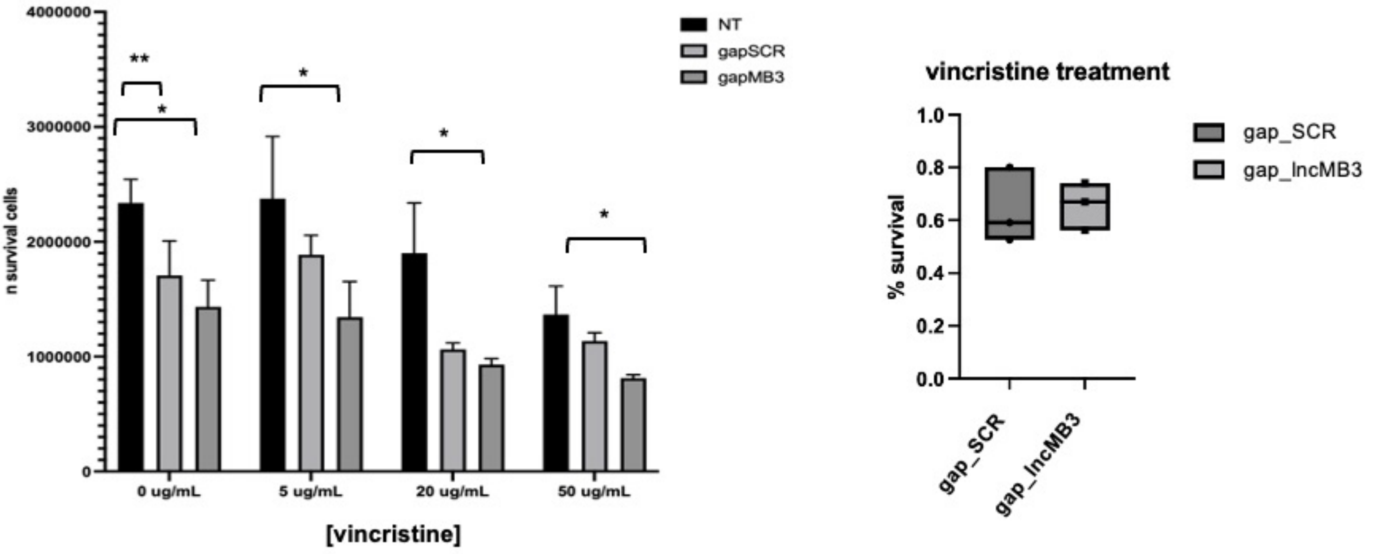


## S8C

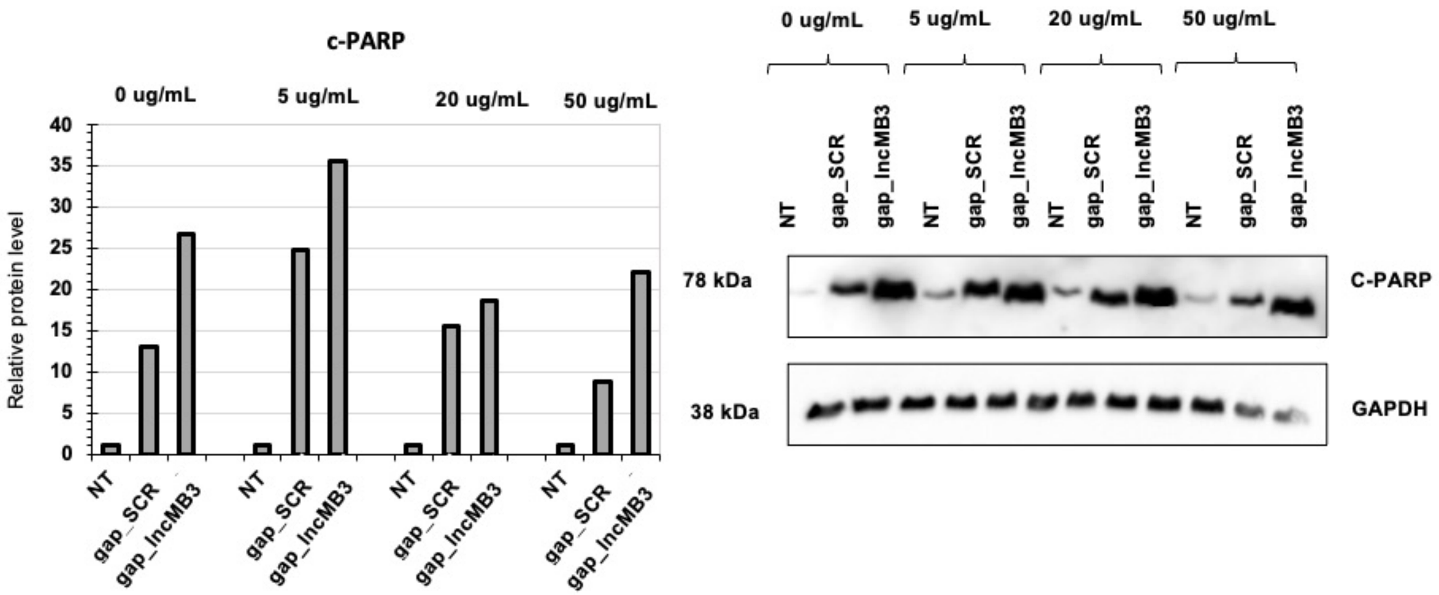

### SUPPLEMENTARY FIGURE 9

#### FIGURE S9

S9A

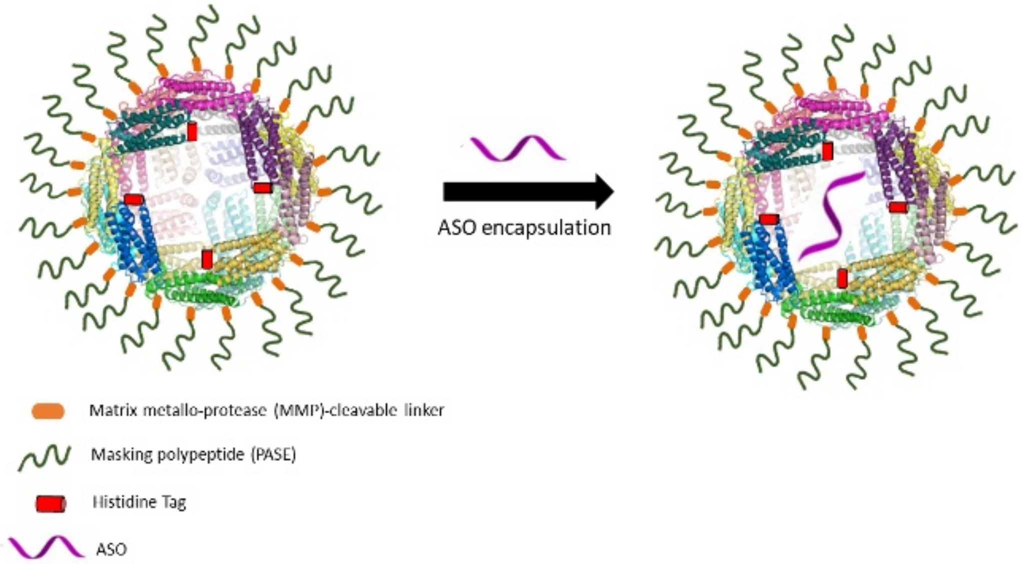


S9B

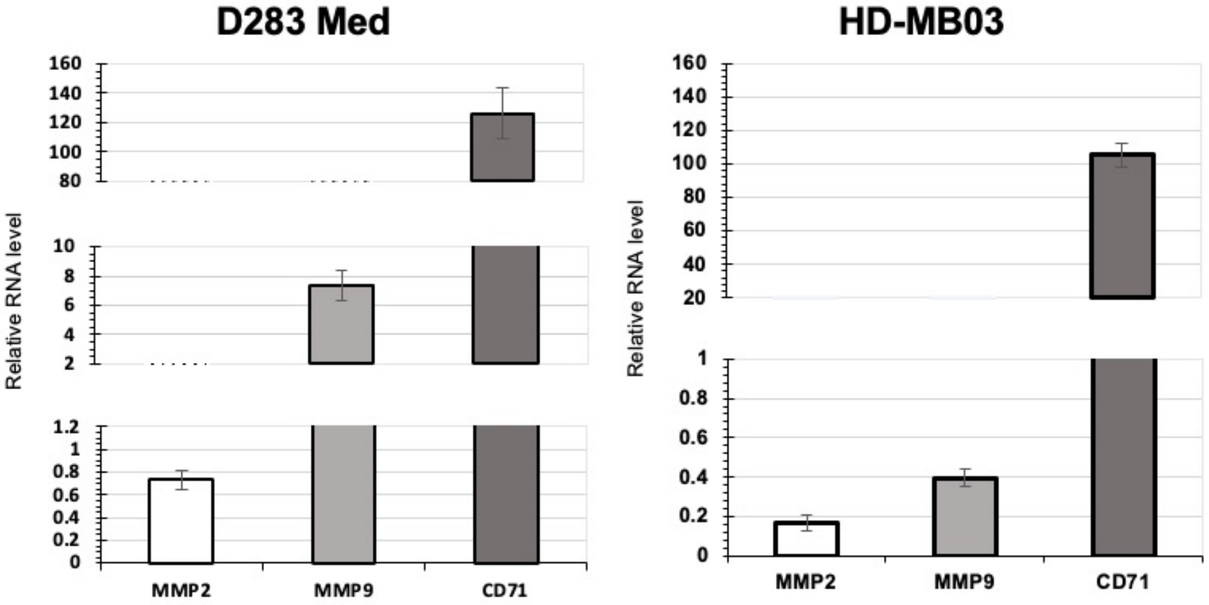


S9C

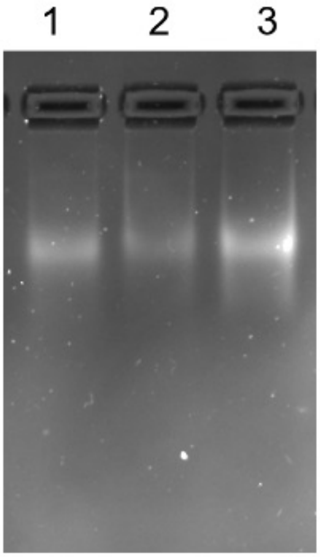


S9D

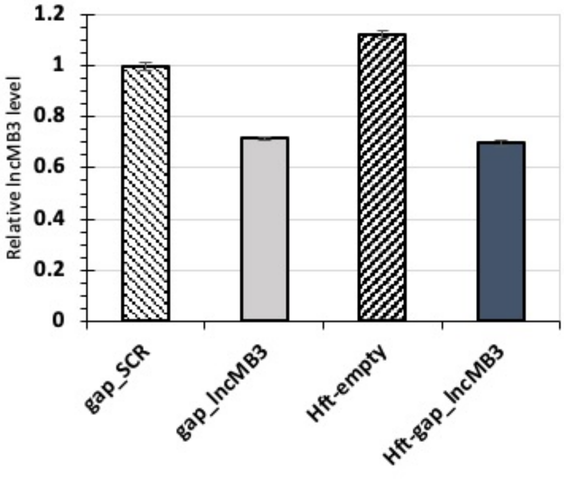
