## SUPPLEMENTARY FIGURE 6 for "A circuit involving the lncRNA MB3 and the driver genes MYC and OTX2 inhibits apoptosis in Group 3 Medulloblastoma by regulating the TGF-β pathway via HMGN5"

### FIGURE S6

## S6A

| Gene | log2FC Even VS inp | FDR even VS inp | log2FC odd VS inp | FDR odd VS inp |
|---|---|---|---|---|
| lncMB3 | 9.523853 | 0.007245 | 7.941424 | 0.028996 |
| HMGN5 | 6.060362 | 0.00112745 | 7.475088 | 0.004114 |
| ANKDD1A | 7.0368 | 0.011791 | 6.388846 | 0.029212 |
| EIF5B | 4.852299 | 0.017849 | 5.803747 | 0.009553 |

## S6B

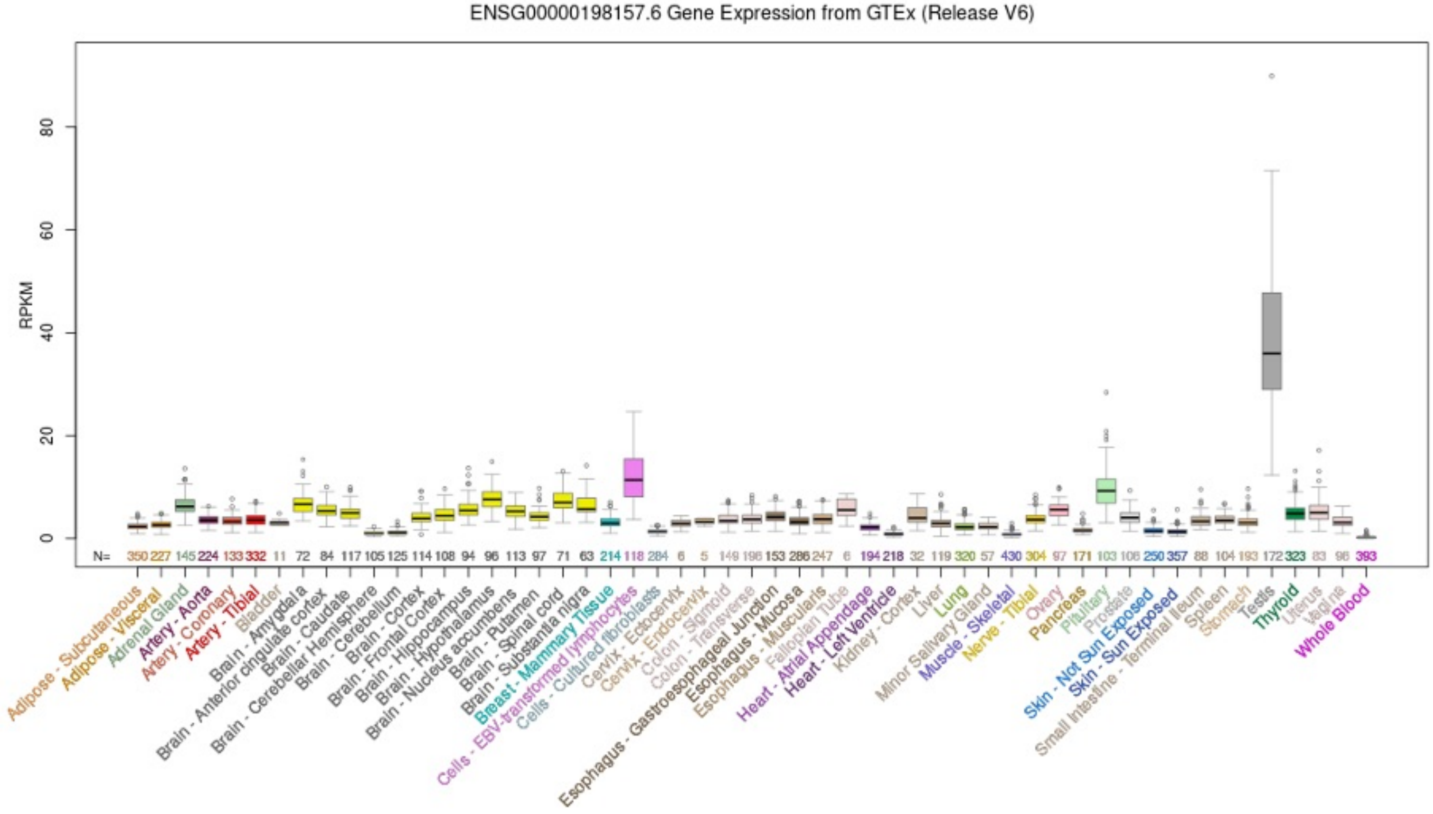


## S6C

| SAMPLE | TARGET | COPY NUMBER |
|---|---|---|
| D283 Med | LncMB3 | 135 |
| D283 Med | HMGN5 | 213.9 |
| D283 Med | GAPDH | 2051 |
