## SUPPLEMENTARY FIGURE 7 for "A circuit involving the lncRNA MB3 and the driver genes MYC and OTX2 inhibits apoptosis in Group 3 Medulloblastoma by regulating the TGF-β pathway via HMGN5"

### FIGURE S7

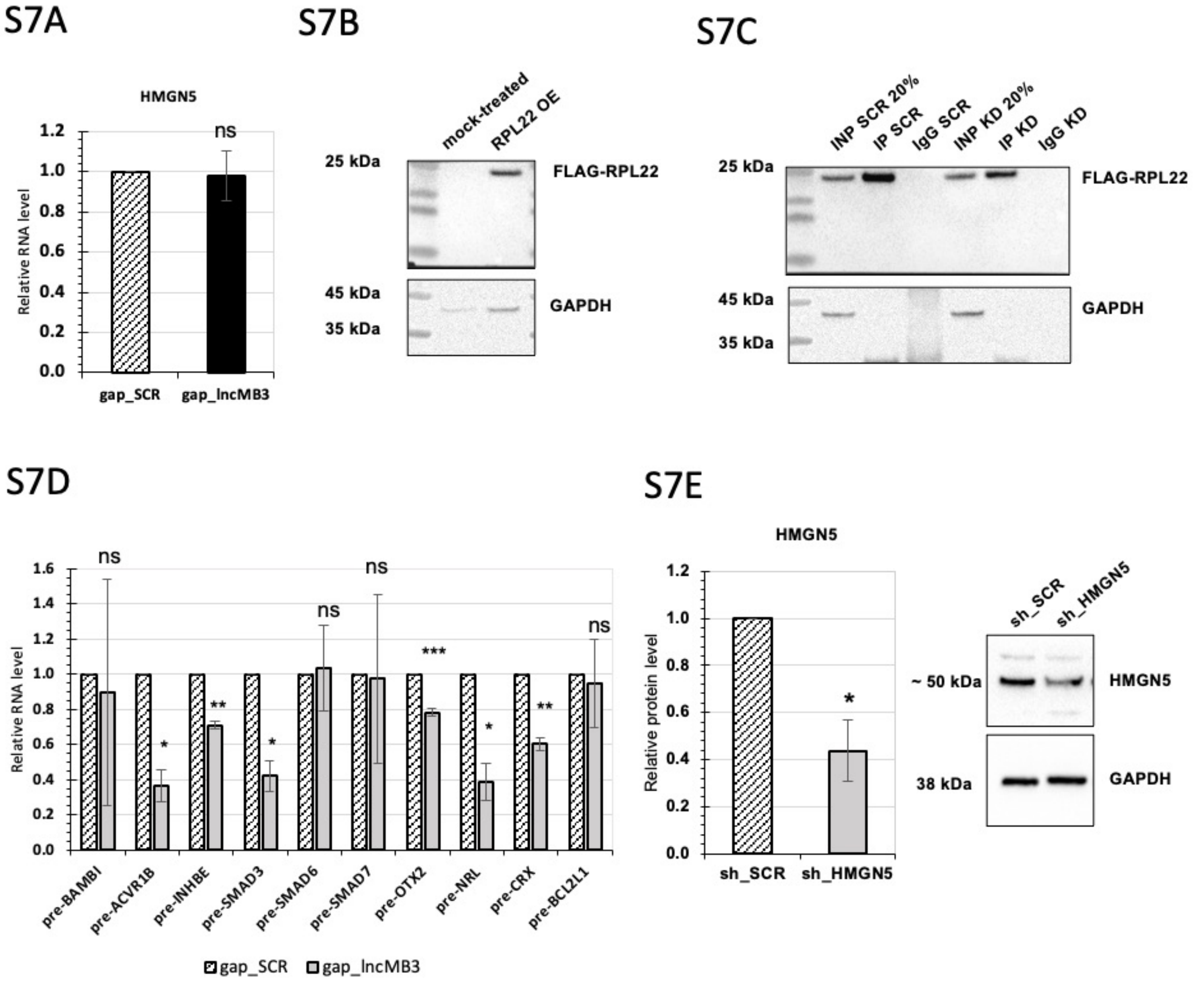


## S7F

| Gene Name | KD | % downregulation |
|---|---|---|
| INHBE | gap_lncMB3 | ~80% |
| | sh_HMGN5 | ~30% |
| SMAD3 | gap_lncMB3 | ~80% |
| | sh_HMGN5 | ~50% |
| OTX2 | gap_lncMB3 | ~80% |
| | sh_HMGN5 | ~20% |
| BCL2L1 | gap_lncMB3 | ~40% |
| | sh_HMGN5 | ~30% |
